# cLoops2: a full-stack comprehensive analytical tool for chromatin interactions

**DOI:** 10.1101/2021.07.20.453068

**Authors:** Yaqiang Cao, Shuai Liu, Gang Ren, Qingsong Tang, Keji Zhao

## Abstract

Investigating chromatin interactions between regulatory regions such as enhancer and promoter elements is vital for a deeper understanding of gene expression regulation. The emerging 3D mapping technologies focusing on enriched signals such as Hi-TrAC/TrAC-looping, compared to Hi-C and variants, reduce the sequencing cost and provide higher interaction resolution for *cis*-regulatory elements. A robust pipeline is needed for the comprehensive interpretation of these data, especially for loop-centric analysis. Therefore, we have developed a new versatile tool named cLoops2 for the full-stack analysis of the 3D chromatin interaction data. cLoops2 consists of core modules for peak-calling, loop-calling, differentially enriched loops calling and loops annotation. Additionally, it also contains multiple modules to carry out interaction resolution estimation, data similarity estimation, features quantification and aggregation analysis, and visualization. cLoops2 with documentation and example data are open source and freely available at GitHub: https://github.com/YaqiangCao/cLoops2.

## Introduction

Chromatin is known to be well organized in multi-scale 3D structures as loops, domains, and compartments (*1, 2*). These structures play important regulatory roles for gene expression (*3, 4*) and biological processes such as development (*5*) and cell cycle (*6, 7*). Transcription factors (TFs), CTCF and cohesin, are major players in establishing chromatin architectures through the loop extrusion model (*8–10*). Other TFs, such as YY1 (*11*), Wapl (*12*), and ZNF143 (*13*), are implicated in 3D dynamic changes of the chromatin. Due to the important roles of chromatin interaction in cells, many efforts have been put forward to developing versatile experimental tools for elucidating the 3D structure of chromatin. Among the popular proximity-ligated based methods, Hi-C unbiasedly detects genome-wide interactions (*14, 15*) while ChIA-PET (*16*), HiChIP (*17*) and PLAC-seq (*18*) detect chromatin interactions over selected genomic regions. Hi-TrAC and TrAC-looping are independent of proximity ligation and detect both chromatin accessibility and interactions at high-resolution (***Liu. et al* in submission, data un-published**) (*19*); therefore, they are useful for analyzing promoter-enhancer interactions. Effective computational pipelines are necessary to intercept the data to draw biological conclusions.

Hi-TrAC and TrAC-looping use the DNA transposase Tn5 to capture chromatin interactions by inserting an oligonucleotide bridge between two interacting chromatin loci. Thus, in addition to providing chromatin interaction information, they also measures chromatin accessibility like ATAC-seq (*20*). Hi-TrAC/TrAC-looping data contain information for interactions between accessibility sites such as enhancers and promoters as loops as well as generic chromatin interactions analogous to Hi-C interactions like domains. Therefore, a comprehensive pipeline to analyze this kind of data should include: 1) preprocessing raw reads into interaction data 2) peak-centric analysis 3) loop-centric analysis 4) domain-centric analysis. The peak-centric analysis methods have been well developed for ChIP-seq data (*21–24*), and multiple domain-centric analysis methods are available for Hi-C data (*25–27*). Loop-centric analysis can reveal regulation details of *cis* regulatory and looping function; however, this category of analysis methods are still underdeveloped. Ideally, loop-centric anlayis tool may consist of core modules of:1) accurate calling algorithms 2) visualization 3) differentially enriched loops calling analogous to differentially expressed genes analysis or sample-wise comparison methods 4) loops annotations 5) integration analysis with other data.

In this study, we introduce a new analysis pipeline, cLoops2, to address the practical analysis requirements, especially for loop-centric analysis with preferential design for Hi-TrAC/TrAC-looping data. Based on an improved highly sensitive, unbiased and unsupervised clustering algorithm blockDBSCAN, cLoops2 directly analyzes the paired-end tags (PETs) to find candidate peaks and loops. It estimates the statistical significance for the peak/loop features with a permuted local background, eliminating the bias introduced from 3^rd^ peak-calling parameters tuning for calling loops. Along with the core modules: peak-calling, loop-calling, differentially enriched loops calling, visualization, features quantification, features aggregation analysis and loops target annotations, cLoops2 also implements many utility functions, for example, estimation of reasonable interactions, which can be determined semi-empirically at some resolution. Although cLoops2 was designed for and applied to analyzing TrAC-looping and Hi-TrAC data, it can also be applied to peak-calling of state-of-art ChIP-seq like sequencing methods and comprehensive loop-centric analysis for ChIA-PET and HiChIP data, showing the versatility of cLoops2.

## Results

### Overview of cLoops2

cLoops2 is a full suite of tools for 3D genomic interaction data, improved from our previous work cLoops (*28*), with many extended modules of functions (**Supplemental Figure 1** and **Figure 1**). Like cLoops, cLoops2 is also a light-weight command line tool for Linux and macOS for use via terminal and Python, which can be easily integrated as a backend component for potentially more sophisticated tools with web or graphical interfaces. The learning curve of cLoops2 is comparable to other tools like SAMtools (*29*), BEDtools (*30*), or MACS2 (*21*). The main modules of cLoops2, such as peak/loop/domain calling algorithms and features aggregation analysis, are integrated into the main command, and can be executed through the prefix cLoops2 (**Supplemental Figure 1**). Meanwhile, extendable supplementary analysis scripts can be executed independently. For example, tracPre2.py processes raw Hi-TrAC (tracPre.py processes TrAC-looping) data from FASTQ files to uniquely high-quality mapped paired-end tags (PETs) BEDPE files (**Supplemental Figure 1**). Each sub-command can be run through the “cLoops2 sub-command” (for example, “cLoops2 callPeaks”), with a program description, parameter details, and examples shown as the default output.

**Figure 1.**
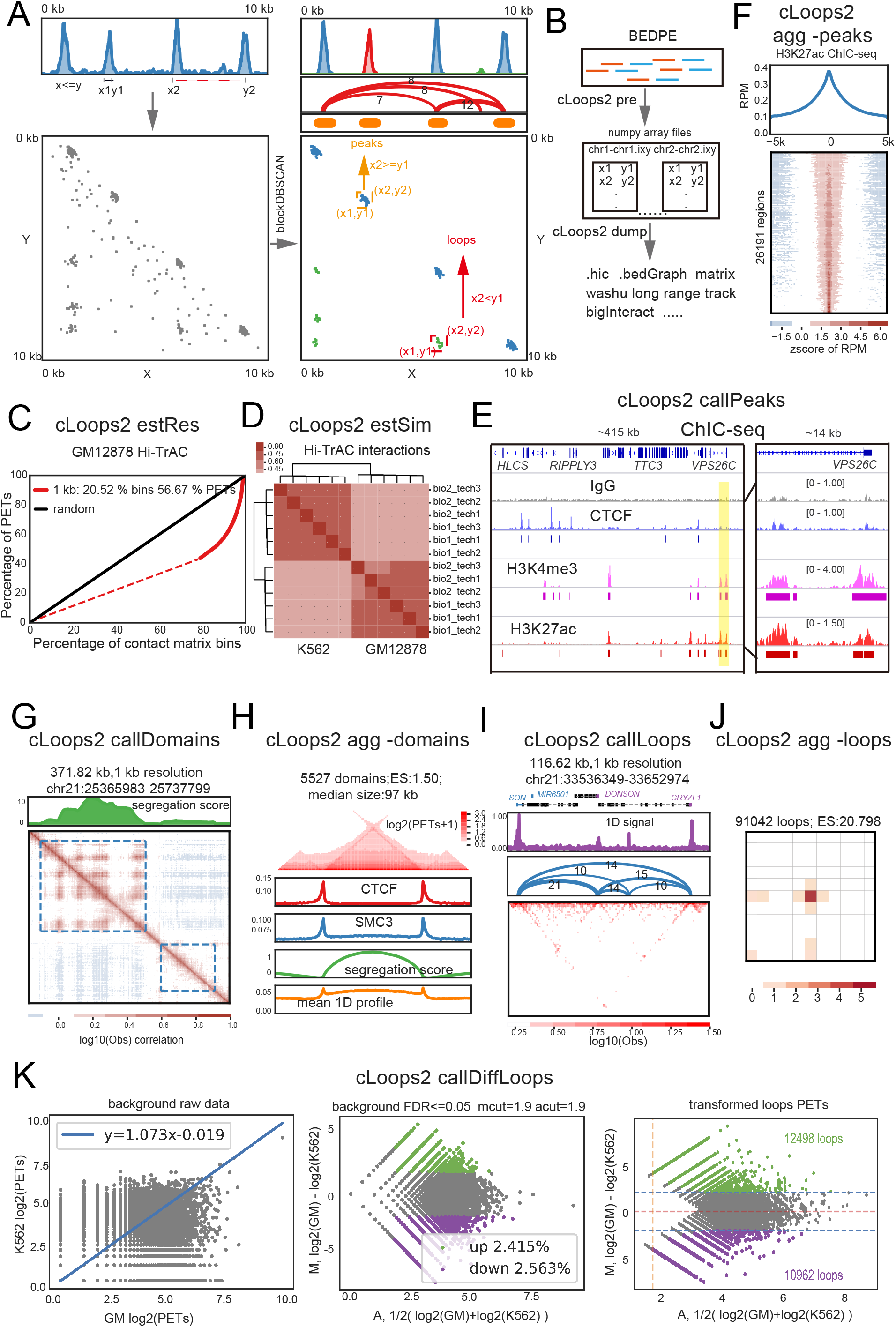
Overview of cLoops2 module features. (A) A demo of cLoops2 peak-calling and loop-calling core algorithms based on blockDBSCAN clustering with hypothetical data. There were peaks, loops, and background noise in the hypothetical data as they mimiced the properties of Hi-TrAC/TrAC-looping or HiChIP data. Processed paired-end tags (PETs) from sequencing data were marked as (x, y) for each one for their mapped coordinates in the genome. When PETs were projected into 2D space, close PETs (such as (x1, y1)) were projected to a diagonal line or nearby, while distal PETs (such as (x2, y2)) were projected away from the diagonal line. After blockDBSCAN clustering, candidate peaks and loops were identified from the clusters. (B) cLoops2 pre and dump modules for data input-output stream. The cLoops2 pre module processes mapped PETs from BEDPE files into cLoops2 specific data files of Numpy array files. cLooops2 dump module converts cLoops2 data to other data types such as HIC file. (C) An example of the cLoops2 estRes module for estimating reasonable contact matrix resolution for GM12878 Hi-TrAC data. The dashed line shows the bins from the contact matrix with only singleton PET, and the solid lines show the bins with multiple PETs. The random line indicates all PETs are evenly distributed in the contact matrix bins. We defined the highest interaction resolution for which there are more than 50% non-singleton PETs in the defined bin sizes (as here for 1kb bin size). (D) An example of the cLoops2 estSim module for estimating interaction similarities with 1kb resolution among Hi-TrAC samples. (E) An example of the cLoops2 callPeaks module for peak-calling of ChIC-seq data. The left panel shows a randomly selected genomic region with the ChIC-seq signals for IgG, CTCF, H3K4me3, and H3K27ac in GM12878 cells and called peaks; the right panel shows the zoom-in region for a better view of captured peaks. (F) An example of the cLoops2 agg module for showing aggregated peaks of GM12878 H3K27ac ChIC-seq data. (G) An example of cLoops2 callDomains module for domain-calling from the GM12878 Hi-TrAC data with key settings of -bs 1000 -ws 100000. Positive segregation scores indicate domain regions. (H) An example of the cLoops2 agg module for showing aggregated domains of GM12878 Hi-TrAC data. (I) An example of the cLoops2 callLoops module for loop-calling from the GM12878 Hi-TrAC data. The cLoops2 plot module generated the plot. (J) An example of the cLoops2 agg module for showing aggregated loops of the GM12878 Hi-TrAC data. (K) Examples of output figures for cLoops2 callDiffLoops module comparing Hi-TrAC loops from GM12878 and K562 cells. Nearby loop background data were used to fit a transformation line between two datasets (left panel). MA plot of background data was used to get the cutoffs for average and fold change with FDR <= 0.05 (middle panel). The transformation line and cutoffs estimated from the background were then applied to loops to find significantly enriched loops (right panel).

The core algorithm for peak-calling and loop-calling in cLoops2 was based on a new clustering algorithm called blockDBSCAN (**Figure 1A**), which further improved cDBSCAN in cLoops. Each PET from the mapping result was marked as (*x*, *y*) for the genome coordinates, with *x* marking the smaller coordinate and *y* marking the bigger one for each end sequencing tag. There are features for both peaks in the 1D linear space and loops in the 2D space with some random noise in both for ideal Hi-TrAC/TrAC-looping, ChIA-PET (*16*), and HiChIP (*17*) data (**Figure 1A**). The PETs with short distance (such as (*x*_1_, *y*_1_) from peaks) in the 1D space were projected to the diagonal line in the 2D space while the long-range interaction PETs (such as (*x*_2_, *y*_2_) from loops) were projected to the space distant from the diagonal line. After blockDBSCAN clustering, noisy points were removed (gray dots in **Figure 1A**), and the number of clusters were automatically determined. Therefore, every cluster can be marked as [(*x*_1_, *x*_2_), (*y*_1_, *y*_2_)], with *x*_1_ marking the minimal coordinate *and x*_2_ marking the maximal coordinate of the left end tags and *y*_1_ marking the minimal coordinate and *y*_2_ marking the maximal coordinate of the right end tags. Based on the boundary overlaps of *x*_2_ and *y*_1_, clusters can be classified into two groups, candidate peaks (*x*_2_ ≥ *y*_1_) and candidate loops (*x*_2_ < *y*_1_). These candidate peaks and loops were tested against the permutated local background to obtain statistical significance to further determine whether they were putative peaks or loops.

### Main modules of cLoops2

The cLoops2 pre module preprocesses the input BEDPE format data of mapped PETs into cLoops2 specific data files as a folder, of which each interacting chromosome pair is saved into a file. All other cLoops2 functions start from this data folder. Each file is a Numpy (*31*) 2D array, recording the mapped coordinates of PETs. cLoops2 dump converts cLoops2 specific data into popular supported file types that can be loaded in Juicebox (*32*) (HIC file), the WashU Epigenome Browser (*33*) (long-range interaction track file), and the contact matrix in a text file that can be loaded into Python, R for further analysis or TreeView (*34*) for visualization (**Figure 1B**). One advantage of the cLoops2 data structure is that the data can be converted back to interacting PETs (cLoops2 dump -bedpe) while other matrix-based data formats such as HIC or cooler (*35*) actually lose the PETs-level information, which limits them to show1D profiles.

The cLoops2 estRes module estimates the resolution of an interaction contact matrix based on singleton PETs and bins (**Figure 1C**). For a specific input resolution, cLoops2 estRes assigns each PET into a contact matrix bin. Accumulation of PETs (Y-axis) against the accumulation bins (X-axis, bins sorted by PETs in the bin ascendingly) for the input resolution was plotted to determine the interaction signal enrichment. If PETs are evenly distributed between genomic locations, there will be a straight diagonal line (black line and red dash line in **Figure 1C**), which is true if we shuffled the two ends of all PETs to an expected background. Otherwise, PETs will be enriched in only a few bins if there are only a few strong interactions. The curve will then show a prominent and steep rise towards the higher ranked bins (red solid line in **Figure 1C**). There are high possibilities that those singleton PET bins are due to noise or background signal. Because of the existence of chromatin domains and loops in the 3D genome, specific PETs should display certain degrees of enrichment. Therefore, we define the highest genome-wide resolution as more than 50% PETs (solid curves in **Figure 1C**) detected in multiple PET bins. The estimation is only a whole genome estimation, and some local regions may have higher or lower resolutions. The cLoops2 estRes module can also estimate similarities among different experimental interaction samples (**Figure 1D**).

There are four main callers for features finding in cLoops2: callPeaks for interaction data such as Hi-TrAC/TrAC-looping data or 1D genome feature profiling data such as ChIC-seq (**Figure 1E**), callDomains (**Figure 1G**), callLoops (**Figure 1I**), and callDiffLoops for calling differentially enriched loops (**Figure 1K**) from Hi-TrAC/TrAC-looping data. For the identified features, different kinds of aggregation analysis were also implemented in the cLoops2 agg module for peaks (**Figure 1F)**, domains (**Figure 1H**), and loops (**Figure 1J**); meanwhile, the cLoops2 quant module can generate similar files to the output of callers if features are called in one sample and quantified in another sample (**Supplemental Figure 1**).

To call differentially enriched loops between two conditions, loops from pairwise samples are combined first and then quantified in both samples. Loops’ nearby regions are defined as background, quantified, and then fitted linearly (**Figure 1K** left panel). FDR is a required parameter, which by default is set to 0.05, to find the cutoffs of average and fold change in an MA plot for background data (**Figure 1K** middle panel). The fitted coefficient and interpret transform the PETs in loops of treatment dataset to control dataset as assuming there should be no difference in background data (**Figure 1K** right panel). The average and fold change cutoffs are then applied to transformed loops data (**Figure 1K** right panel). Poisson *P*-value is assigned to each loop as 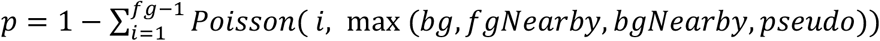. Where fg stands for the bigger value of PETs in the testing loop for treatment against the control sample; bg stands for the smaller value for comparison; fgNearby is the number of PETs from background data for the testing condition, and bgNearby is the number of PETs for background data from the other condition. Pseudo is a general noise control value set to 1 as a default and can be adjusted through its parameter. The Poisson *P*-values are further corrected by the Bonferroni correction.

### The blockDBSCAN algorithm

In a previous study of cLoops, we proposed cDBSCAN, which was improved from the popular DBSCAN algorithm, for clustering interacting PETs directly to find candidate loops (*28*). Analyzing PETs directly achieves resolution scale-free performance, which is an important feature to identify loops’ variable sizes and accurate boundaries. Although cDBSCAN worked well for previously tested ChIA-PET, TrAC-looping, HiChIP, and Hi-C data, it is quite time-consuming for deeply sequenced libraries (*36*). Therefore, we further improved the algorithm’s speed and named it blockDBSCAN (**Figure 2**). In cDBSCAN, after indexing and noise removal, clustering is performed comparable to DBSCAN for all remaining PETs but with a limited search to neighbor index blocks. In the last step, each index block can be treated as a point with weight, with the coordinate being the geometric mean of PETs, and the weight being the number of PETs in that index block (marked from a to f in **Figure 2A**). Clustering performed at the level of index blocks can effectively reduce algorithm complexity, and therefore increase clustering speed. We evaluated the performance of blockDBSCAN against cDBSCAN with simulated data, and indeed confirmed that blockDBSCAN was faster and even a little better matched with true simulated labels of signals measured by the adjusting rand index (ARS) (**Figure 1B**). We further validated the clustering algorithm efficacy for finding candidate loops with real Hi-TrAC data, CTCF ChIA-PET data, Hi-C data, and H3K27ac HiChIP data (**Figure 2C**), achieving about 2 to 15 folds speed improvement and with nearly the same memory usage (**Supplemental Figure 2**). blockDBSCAN was estimated approximate *O*(*N*) complexity as run time increases nearly linearly with smaller slopes comparing to cDBSCAN when the number of PETs increases (**Figure 2C**).

**Figure 2.**
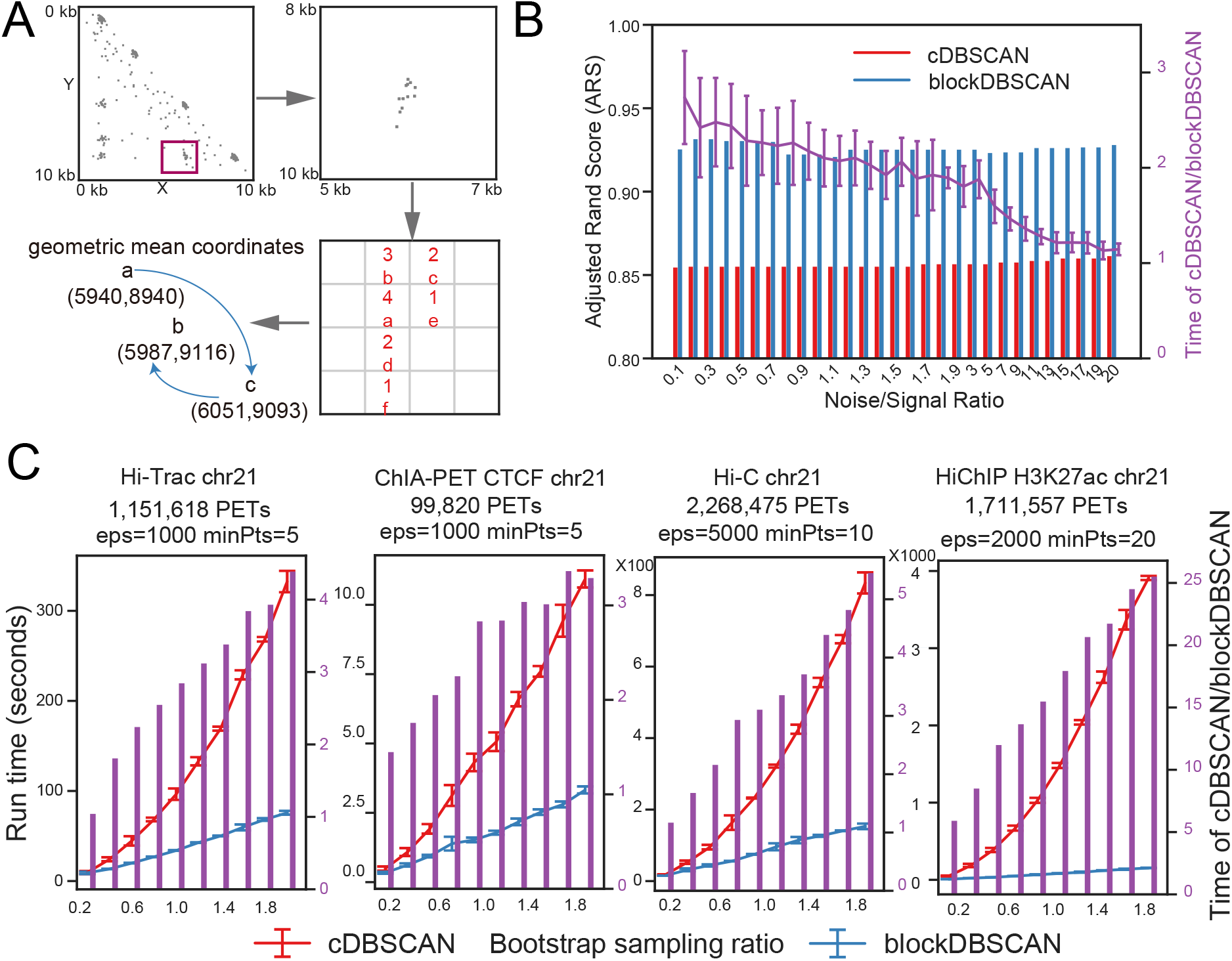
The blockDBSCAN algorithm. (A) A demo of blockDBSCAN algorithm processes on hypothetical data. The red rectangle highlights the region for detailed demonstration. After indexing and noise removal, informative PETs were located in the indexed blocks with the side length of eps (one key parameter of DBSCAN). For each indexed block, marked as a to f, PET numbers in them were also annotated. Geometric mean coordinates were computed for each indexed block from PETs in that indexed block for the following clustering at the block level. (B) The comparison of running CPU time and accuracy of cDBSCAN and blockDBSCAN at different noise/signal ratios was based on 5 repeats of simulation data. The left y-axis marked the bars for adjusted rand scores (ARS), showing consistency with simulation data true labels; the right y-axis marked the error bars for running time ratios. ARS measured the similarity between clustering results and true labels ranging from −1.0 to 1.0, with 0 indicating random labeling and 1 is a perfect match. The simulation data were generated in the same way described in cLoops (*28*). (C) Comparison of running CPU time of cDBSCAN and blockDBSCAN with real interaction data of chromosome 21 from Hi-TrAC, CTCF ChIA-PET, in situ Hi-C, and H3K27ac HiChIP data in GM12878 cells (**Supplemental Information**), based on 5 repeats and sampling.

### Peak-calling for next-generation ChIP-seq like data

Peak-calling is a fundamental analysis step for popular 1D genomic feature profiling methods such as ChIP-seq (*37*) and ATAC-seq (*20*). Currently, the most popular peak-calling algorithms were designed for ChIP-seq, such as MACS (*21*) but have limitations such as reduced precision (*38*) for the latest emerging technologies such as CUT&RUN (*39*) and ChIC-seq (*40*). A possible explanation for that is a different signal-to-noise ratio and variable peak sizes compared to ChIP-seq. Loop-calling tools, such as Mango (*41*) for ChIA-PET data and hichipper (*42*) for HiChIP data, usually start from peak-calling, and then combinations of peaks are used to find candidate loops. It is inferable that inaccurate peak-calling may lead to false positive candidate loops for those tools. Meanwhile, it was also noticed the peak-calling results from both FitHiChIP (*43*) and hichipper (*42*) were strongly biased due to HiChIP specific biases (*44*), which may bias the accuracy of finally called loops. Thus, to date, generalist peak-calling algorithms are still needed. In a previous study of cLoops (*28*), we only used cDBSCAN to identify candidate loops. Now we found that the density approaching principle can also be applied to identify candidate peaks (**Figure 1A**).

To benchmark the performance of the cLoops2 callPeaks module in calling peaks, we generated ChIC-seq datasets for the transcription factor CTCF with sharp peaks (**Figure 3A**), the histone modification H3K4me3 with sharp peaks (**Supplemental Figure 3A**), the histone modification H3K27ac with both sharp and board peaks (**Figure 3B**), and IgG ChIC-seq control data. The peaks called from these datasets were compared with three popular ChIP-seq peak callers: MACS2 (*21*), SICER (*45*), HOMER (*46*), and SEACR (*38*) which was designed for CUT&RUN protocol (**Methods)**.

**Figure 3.**
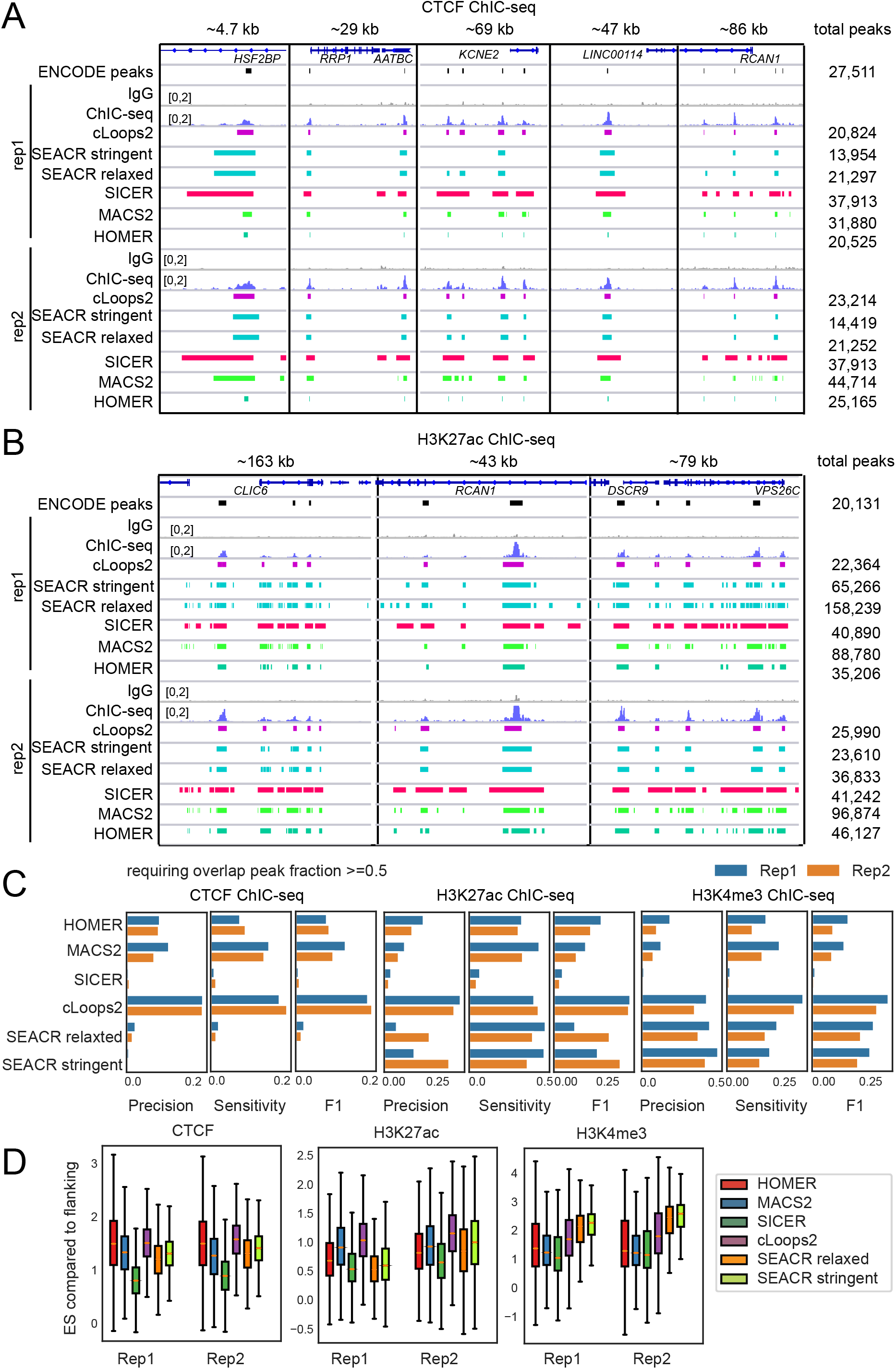
Peak calling by cLoops2 from ChIC-seq data. (A) Genome Browser images showing the CTCF ChIC-seq data and peaks called by different tools. Two biological replicates of CTCF and IgG ChIC-seq datasets were shown. (B) Genome Browser images showing the H3K27ac ChIC-seq data and peaks called by different tools. (C) Precision, sensitivity, and F1 scores for peaks called by different tools by comparing with the ENCODE peaks (**Metthods**), requiring peaks overlap fractions >=0.5. (D) ChIC-seq signal-to-noise enrichment score (ES) from various tools. The ES was calculated by comparing signals in the peak region with that in the flanking upstream and downstream same-sized regions.

We first examined the examples of called peaks from the CTCF ChIC-seq data and compared them to the ENCODE reference peaks from ChIP-seq (**Supplemental Information**, **Methods**). We noticed that cLoops2 captured most of the major peaks in three randomly chosen regions, which were well-aligned with the ENCODE reference peaks (**Figure 3A**). The density-based approach also found accurate boundaries of peaks (**Figure 3A**). The peaks called by both SEACR’s stringent and relaxed modes were too broad, and the SEACR stringent mode missed some significant peaks, resulting in a limited total number of peaks found (**Figure 3A**). SICER was designed for broad histone modification peaks, and as expected, peaks were too broad for transcription factor peaks such as CTCF binding sites (**Figure 3A**). MACS2 yielded the most peaks in replicate 2 (44,714); however, many of them were most likely background regions located near true peaks (**Figure 3A**). Meanwhile, for MACS2, fewer peaks were called for replicate 1 (31,880); the difference in numbers of peaks in the two replicates resulted in lower consistency between replicates (**Figure 3A** and **Supplemental Figure 3D**). The high sensitivity of MACS2 may explain this. HOMER achieved similar performance as cLoops2 for CTCF results (**Figure 3A**). Compared to the cLoops2 results, peaks called by HOMER were sharper, which were apparently much narrower than the ChIC-seq peaks as shown by the Genome Browse images, indicating improper capturing of peak boundaries.

Regarding the peak calling from the H3K4me3 ChIC-seq data, cLoops2 also worked well (**Supplemental Figure 3A**). Both the SEACR stringent and relaxed modes missed some significant peaks, resulting in much lower numbers of peaks as compared to the ENCODE peaks (**Supplemental Figure 3A**). The default SICER setting resulted in some peaks stitched together (**Supplemental Figure 3A)**. MACS2 called the most peaks for both replicates; again, many of them may be just background signals near true peaks. HOMER achieved similar performance with cLoops2 for calling H3K4me3 peaks (**Supplemental Figure 3A**).

H3K27ac displays both sharp and broad peaks in the genome (**Figure 3B**). cLoops2 detected most of the significant reference peaks, called similar peaks as reference peaks, and showed high consistency between two biological replicates (**Figure 3B**). Both SEACR stringent and relaxed mode called too many peaks and were too sensitive for replicate 1 (**Figure 3B**). Similar to SEACR, SICER was too sensitive and called too many broad peaks (**Figure 3B**). MACS2 and HOMER called many background regions near highly enriched peaks (**Figure 3B**).

Precision, sensitivity, and the F1 score (*47*) were used as metrics to evaluate the performance of the genome-wide peak-calling by comparing the discovered peaks with the ENCODE reference peaks (**Methods**). If requiring a minimal overlap of 1 bp between called peaks, then broad peaks had overestimation of precision and sensitivity (**Supplemental Figure 3B**) in the situation that only small parts of the called peaks overlapped with reference peaks at a low overlapping ratio. Therefore, we required half of the candidate peaks to overlap with the reference peaks to get evaluation metrics. With this requirement, cLoops2 had the highest F1 scores for all the three ChIC-seq datasets, showing both high precision and sensitivity for calling accurate peaks (**Figure 3C**). cLoops2 also showed comparable reproducibility between replicates (**Supplemental Figure 3C**) and correlation between gene expression and the peak density at the transcription start sites (**Supplemental Figure 3D**). Assuming signals from same region of IgG control of a putative peak and the putative peak’s flanking regions are background, then we can use the signal-to-noise ratio distribution to check whether putative peaks are properly called. Missclassification of background regions as putative peaks or vice versa will both decrease the distribution of signal-to-noise ratios. Further comparison of signal densities in peaks against their flanking upstream and downstream regions of the same size (**Figure 3D**) and against the same regions in the IgG control sample (**Supplemental Figure 3E**), especially for CTCF and H3K27ac, indicates that cLoops2 performed the best to capture peaks with proper boundaries.

cLoops2 also demonstrated excellent performance for peak calling from CTCF and H3K27ac CUT&RUN data, as shown both by the Genome Browser snapshots of random selected examples (**Supplemental Figure 4A** and **B**) and the whole genome evaluation metrics (**Supplemental Figure 5**). Due to the differences between the ENCODE reference peaks of H3K27me3 and CUT&RUN data, it is currently a challenge to carry out a fair comparison of peak-calling performance for H3K27me3 (**Supplemental Figure 4C**). cLoops2 also achieved the best performance for H2A.Z, H3K4me1, and H3K27ac CUT&TAG data measured by F1 score compared to the corresponding ENCODE reference peaks (**Supplemental Figure 6**). It should be noted that SEACR was designed for CUT&RUN series technology with low background, while cLoops2 was designed as a general peak-caller based on the density approaching principle. However, even for the CUT&RUN series data, cLoops2 works well and, to some extent, better than SEACR, indicating that the principles of the cLoops2 peak-calling algorithm can be applied to potentially more genomic features profiling technologies under development.

**Figure 4.**
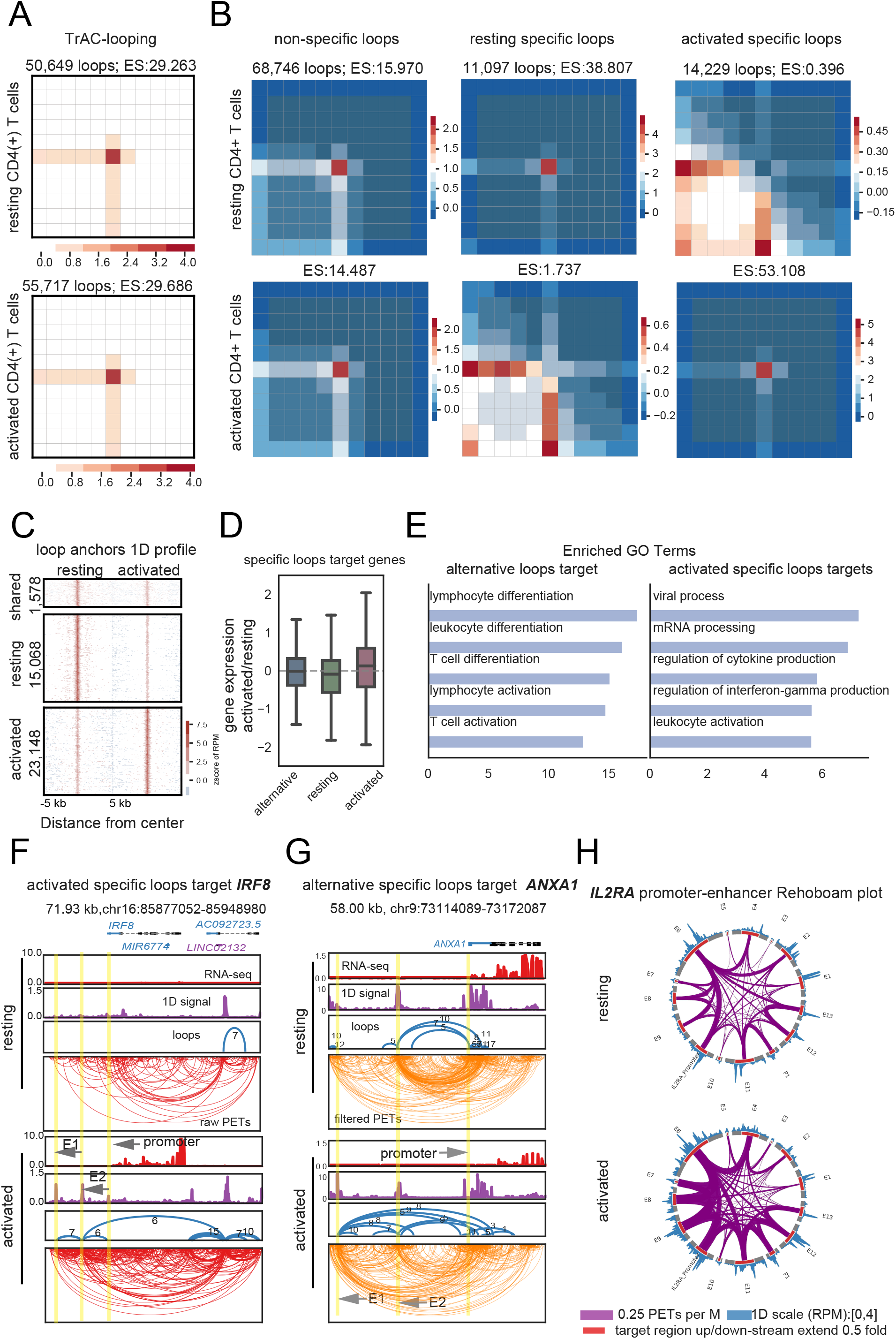
cLoops2 calls differentially enriched loops from TrAC-looping data. (A) Aggregation analysis of called loops from TrAC-looping data of resting and activated CD4+ T cells. Mapped cis-PETs were sub-sampled to 100 million for analysis and comparison. Enrichment score (ES) was the average value of the matrix center divided by others of all loops. (B) Aggregation analysis of non-specific and specific loops between resting and activated CD4+ T cells. The cLoops2 callDiffLoops carried out the analysis. (C) Aggregation analysis of TrAC-looping 1D signals on combined anchors from cell-specific loops. Only limited anchors were shared between different cells. The specific 1D signals also indicated cell-specific accessibilities of the anchors. (D) Gene expression associations with cell-specific loops. Alternative means a gene’s promoter is looping by different enhancers in two conditions. (E) Enriched gene ontology (GO) terms for cell-specific loops targeting genes. Only the top 5 were shown. (F) Example of activated T cell specific loops targeting gene *IRF8*. Blue color marks the first exon of the gene in the positive strand, and purple color marks the first exon of the gene in the negative strand. E1 and E2 mark the enhancers that were active only in the activated CD4 T cells. (G) Example of alternative specific loops targeting gene *ANXA1*. Only PETs overlapped with any end of loop anchors were remained, performed with the cLoops2 filterPETs module. (H) Example of Rehoboam plots of interacting enhancers and promoters of the *IL2RA* gene locus. Putative enhancers were marked as E1 to E13, inferred from loop anchors interacting with I*L2RA*’s promoter from the two conditions (**Supplemental Figure 8**). A Rehoboam plot can be obtained through the cLoops2 montage module. TrAC-looping 1D profiles were shown outside the circle, and the levels of interaction were shown as widths of arches that link the interacting regions.

**Figure 5.**
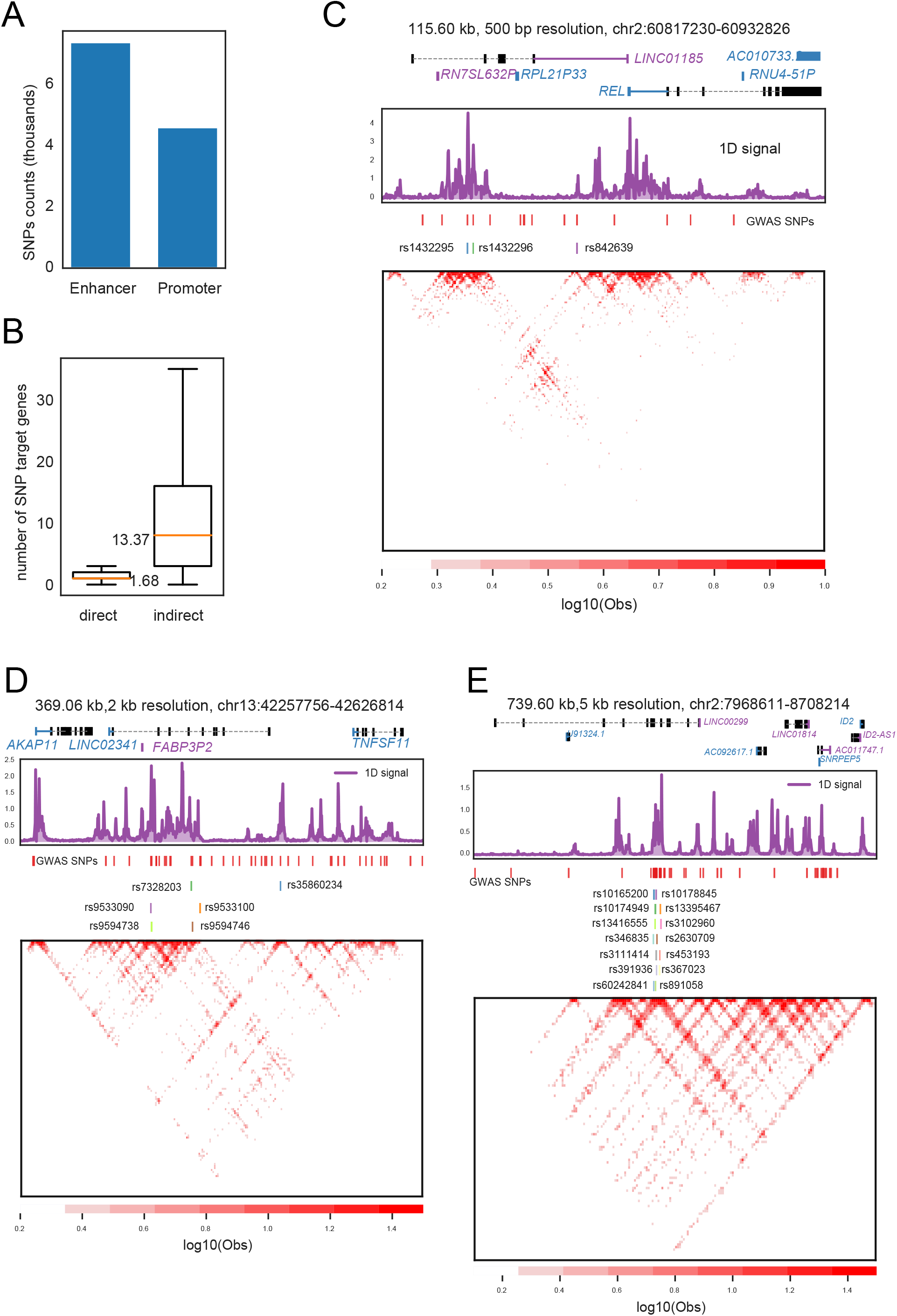
cLoops2 identifies SNP target genes through loops. (A) A large number of the GWAS catalogged SNPs overlaped with the Hi-TrAC loop anchors in GM12878 cells. (B) The direct and indirect target genes of GWAS catalogged SNPs. The target genes were annotated by cLoops2 with loops called from GM12878 Hi-TrAC data. A direct target gene refered to a gene that an SNP was located in one of the loop anchors and the other anchor of the loop was located to the promoter region of the gene. An indirect target gene refered to a gene that an SNP was located in a loop anchor and the other anchor of the loop was not located directly to the promoter region of the gene but instead was linked to the promoter through one or more loops. (C) Example of SNP target gene identified by Hi-TrAC loops. The SNPs were mapped to the lincRNA gene *LINC01185*, while the Hi-TrAC data indicated that the SNPs actually targeting the *REL* gene through enhancer-promoter looping. Blue color marked the first exon of the gene in the positive strand, and purple color marked the gene’s first exon in the negative strand. SNPs colored other than red were SNPs found located in lncRNA but were looped to the promoter regions of other genes. (D) Example of SNP target gene identified by Hi-TrAC loops. The SNPs were mapped to the lincRNA gene *LINC02341*, while the Hi-TrAC data indicated that the SNPs actually targeting the *AKAP11* and *TNFSF11* genes through chromatin looping. (E) Example of SNP target gene identified by Hi-TrAC loops. The SNPs were mapped to the lincRNA gene *LINC00299*, while the Hi-TrAC data indicated that the SNPs actually targeting the *ID2* gene through chromatin looping.

### Comprehensive analysis of loops for interaction data

Many tools have been proposed for calling loops for Hi-C (such as HiCCUPS (*48*), SIP (*36*), and Mustache (*35*)) and HiChIP (such as hichipper (*42*) and FitHiChIP (*43*)). Most of them were specifically designed for only one limited data type, restricted to resolution settings and lacking downstream analysis modules. To our knowledge, cLoops is still the only tool whose principle can be applied to a broad range of interaction data and is not limited by specific resolutions unsupervisely. Therefore, we extended the core of the loop-calling algorithm in cLoops to cLoops2 with more loop-centric analysis.

TrAC-looping data (*19*) were used to benchmark the performance of cLoops2 in loop-calling using the cLoops2 callLoops module and to identify differential loops between different samples using the cLoops2 callDiffLoops module. As blockDBSCAN is more sensitive than cDBSCAN, cLoops2 was able to detect more loops than cLoops (33,416 loops called by cLoops and 50,649 loops called by cLoops2, respectively, from the resting TrAC-looping data). While the loop anchors called by cLoops were not examined for any peak properties, loop anchors called by cLoops2 were significant peaks compared to the nearby regions by default. Therefore, in the aggregation plot, there were continuous signals from the center to the edges (**Figure 4A**). As exemplified with a 17.8kb genomic region, which contained the *IRF2BP2* gene, cLoops2 detected complex and different looping structures between resting and activated CD4+ T cells (**Supplemental Figure 7A**). Markedly more loops were identified around the *IL2* gene, which is a critical gene required for T cell proliferation (*49*), in the activated CD4+ T cells than in the resting CD4+ T cells (**Supplemental Figure 7B**), indicating that cLoops2 can identify dynamic loops associated with cellular functions. Aggregation analysis of cell-specific loops and their nearby regions revealed 11,097 highly enriched loops in resting CD4+ cells and 14,229 in activated cells, respectively (**Figure 4B**). Both sets of the specific loops exhibited a lower or similar level of signals than the nearby regions in the other cell type **(Figure 4B**). For these cell-specific loops, we further examined their anchors of TrAC-looping 1D signals, and found only a few of the anchors (1,578) were shared by the two cell types while most of them were cell-specific (15,068 in resting CD4+ cells and 23,148 in activated CD4+ cells) (**Figure 4C**). The observation that most anchors of the cell-specific loops were also cell-specific suggests two interesting possibilities: 1) cell-specific accessibility was guided by cell-specific looping structures 2) cell-specific looping was guided by some key TFs such as pioneer factors, which can be detected by 1D profiling methods such as ChIP-seq, DNase-seq and ATAC-seq.

We further annotated these cell-specific loops by the cLoops2 anaLoops module to identify their target genes and examine the target gene expression (**Figure 4D**). If a gene promoter looped with multiple enhancers and was associated with distinct loops in different cell types, it was termed an alternative target. Otherwise, if a gene promoter had loops only in one cell type, it was defined as either resting or activated cell-specific target gene. Globally, resting or activated CD4+ T cell-specific loops targeted specific genes had a little higher expression, but not significantly, in the corresponding condition (**Figure 4D**). For the alternative target genes, even though they did not show significant global expression differences between the two cell types, many of them used different enhancers between resting CD4 and activated T cells, which were enriched in lymphocyte differentiation and activation (**Figure 4E**). The *IRF8* gene is a critical regulator of immune function. Its expression was not detectible and no significant loops of the *IRF8* promoter were identified in the resting CD4+ T cells (**Figure 4F**). In the activated CD4+ T cells, the *IRF8* gene locus exhibited complex looping structures including both promoter-enhancer and enhancer-enhancer loops, accompanied by high levels of *IRF8* expression (**Figure 4F**). E1 and E2 were looped together; E2 was looped with the promoters of both the *IRF8* and *AC092723.5* genes. This example showed clear evidences of gene expression regulation by enhancer-promoter loops upon T cell activation. On the other hand, the regulation of alternative looping target genes might be much more complex and need further study. For example, the *ANXA1* gene was expressed at a similar level in the resting and activated CD4+ T cells (**Figure 4G**). Multiple enhancers were looped with the *ANXA1* gene promoter in the resting and activated CD4 T cells. While E1 had stronger interaction signals with the *ANXA1* promoter in the activated CD4+ T cells, E2 showed stronger interactions in the resting CD4+ T cells. This example demonstrated that even for the non-differentially expressed genes in different cell types, the regulatory mechanisms by enhancers might be different in different cells.

The quantitative measurement of interactions among transcriptional regulatory elements such as enhancers and promoters is important to the mechanisms of gene expression regulation. However, the current interaction heatmaps do not provide such quantitative information and reveal the fine looping structures, especially when comparing different cell types or conditions. For example, the *IL2RA* gene promoter interacted with a total of 13 potential enhancer regions in the resting and activated CD4+ T cells together. It was difficult to see quantitative changes from the interaction heatmaps (**Supplemental Figure 8**). Although the arch loop plots showed overall changes of interaction patterns at this genomic locus, it was difficult to directly visualize the dynamic changes of promoter interactions (**Supplemental Figure 8**). To facilitate a direct visualization of changes in promoter-enhancer interactions, we introduced the Rehoboam plots (**Figure 4H**). In a Rehoboam plot, each target region with extended nearby regions was shown as a part of a circle, TrAC-looping 1D profiles were shown outside the circle, and interaction signal levels were shown as the widths of arches between interacting regions. With the Rehoboam plots, it was easy to conclude that (1) globally more interactions were observed with the *IL2RA* promoter in activated CD4+ T cells; (2) E6 and E11 were looped together in activated CD4+ T cells but not in resting CD4+ T cells; and (3) E9 and E12 had much higher accessibility in activated CD4+ T cells (**Figure 4H**). Thus, a Rehoboam plot served as a useful tool to visualize dynamic changes of interactions and accessibilities of regulatory regions for a specific locus of interest.

We also demonstrated the performance of loop calling by cLoops2 from the latest RAD21 ChIA-PET data (*50*) (**Supplemental Figure 9**) and H3K27ac HiChIP data (*51*) (**Supplemental Figure 10**). The anchors from both the RAD21 specific loops and H3K27ac specific loops showed high cell-type specificity (**Supplemental Figure 9C** and **Supplemental Figure 10C**). Higher gene expression levels were also observed for those target-specific loops (**Supplemental Figure 9D** and **Supplemental Figure 10D**). As RAD21 or H3K27ac guides these loops, we also observed a high binding difference from the ChIP-seq data at loop anchors (**Supplemental Figure 9E** and **Supplemental Figure 10E**). These results indicated that cLoops2 can be applied to different datasets from TrAC-looping, Hi-TrAC, to ChIA-PET, and HiChIP.

### GWAS SNPs target annotation from loops

It is recognized for a long time that most of the disease or trait-associated single-nucleotide polymorphisms (SNPs) based on the genome-wide association studies (GWAS) are in non-coding regions in the human genome (*52, 53*). Transferring the associations to functional causality requires accurately annotating the target genes for the SNPs in the current post-GWAS era (*54, 55*). Since cLoops2 is capable of annotating enhancers to their target genes from TrAC-looping or Hi-TrAC data, it can facilitate the annotation of target genes for the SNPs located with loop anchors. For a total of 116,411 SNPs annotated in the NHGRI-EBI GWAS Catalog (*53*), 11,995 (10.3%) of them were overlapped with the loop anchors identified in GM12878 cells and more than 6,000 of them were located in the enhancer regions of loop anchors (**Figure 5A**). Overall, for the SNPs mapped to the loop anchors, a SNP had an average of 1.68 target genes, and had 3.37 indirect target genes in the loop network by average (**Figure 5B**). In NHGRI-EBI GWAS Catalog annotations, we noticed that if a SNP was located in long non-coding (lncRNA), its target gene was annotated as lncRNA. However, we found that many lncRNAs were located in loop anchors and thus they may regulate target gene expression by chromatin looping. We showed here three examples of lncRNA genes hosting SNPs, which were looped to promoters or enhancers detected by Hi-TrAC (**Figure 5C, D** and **E**) but not shown from GM12878 RAD21 ChIA-PET data (*50*), H3K27ac HiChIP data (*51*), or Hi-C (*15*) data (**Supplemental Figure 11**). Thus, these SNPs might regulate target gene expression through direct enhancer-promoter loops. One single SNP might also regulate multiple distant genes from these annotations. For example, rs7328203, rs9533100, rs35860234, and rs9594746, interact with both the *AKAP11* and *TNFSF11* promoters (**Figure 5D**), implying potentially that these SNPs regulated the expression of both genes.

## Conclusion and Discussion

To achieve genome-wide high resolution of chromatin interaction maps such as 1kb and higher resolution, Hi-C (final 3.7 billion PETs) (*15*), its derivatives CAP-C (final 3.2 billion PETs for CAPC-BLCAPC_B01, other samples all nearly more than 1 billion PETs) (*56*) and Micro-C (final billion PETs) (*57*) all need exhaustive sequencing. The huge sequencing depth requirement limits the method application for more samples and puts a great burden on analyzing them both in time and in memory. Meanwhile, interaction enrichment methods such Hi-TrAC represents another profiling perspective: obtaining comprehensive interactions among accessibility sites with reasonable sequencing depth (100 million informative PETs for all downstream analysis). We present cLoops2 here for comprehensive interpretations of Hi-TrAC and similar data, especially for loop-centric analysis.

In conclusion, cLoops2 showed the general applicability of the density-based clustering algorithm blockDBSCAN for finding features in sequencing-based enriched data derived from either ChIP-seq like 1D profiling technologies or interaction data loop-centric analysis. We anticipate that cLoops2 will be a valuable and popular tool for the broader research community, especially for similar data to Hi-TrAC.

Large scales samples of ChIP-seq have been studied by ENCODE (*58*) and NIH Roadmap Epigenomics (*59*), and large scales samples of interaction data led by 4DN (*60*) consortia and other studies are emerging (*50*). More analysis challenges will be revealed with the accumulation of interaction data, and we anticipate the popularity of cLoops2 for the community.

## Methods

### ChIC-seq experiments

Small cell number ChIC-seq assays were performed similarly to our previous study (*61*). Briefly, GM12878 cells were fixed for 10 minutes in 1% formaldehyde in the DMEM medium. The reaction was stopped by adding 0.125 M glycine. Following permeabilization of 200,000 fixed GM12878 cells with 600 μl RIPA buffer with 0.1% SDS at room temperature for 5 minutes, the cells were rinsed with 600 μl binding buffer twice (10 mM Tris-Cl, 1 mM EDTA, 150 mM NaCl, 0.1% TX-100) and re-suspended in 250 μl binding buffer. Before mixing with the cells, antibodies ((CTCF (Millipore, 47-729), H3K4me3 (Millipore, CS200580), H3K27ac (Abcam, ab4729), IgG (Millipore, NI03)) and PA-MNase were pre-incubated with a molecular ratio of 1:2 at 4 °C for 30 minutes in 50 μl binding buffer for generating antibody+PA-MNase complexes. Then, the permeabilizated GM12878 cells in 50 μl binding buffer were mixed with each antibody+PA-MNase complex and incubated for 1 hour at 4 °C. Cells were washed using 600 μl RIPA buffer (10 mM Tris-Cl, 1 mM EDTA, 150 mM NaCl, 0.1% SDS, 0.1% NaDOC, 1% Triton X-100) four times and pelleted by centrifugation at 900g for 1 minute. Next, cells were rinsed using 200 μl rinsing buffer (10 mM Tris-Cl and 10 mM NaCl, 0.1% TX-100). MNase digestion reactions were activated by re-suspending cells in 40 μl reaction buffer (10 mM Tris-Cl, 10 mM NaCl, 0.1% TX-100, 2 mM CaCl_2_) and incubated at 37 °C for 3 minutes. Reactions were stopped by adding 80 μl stop buffer (20 mM Tris-Cl, 10 mM EGTA, 20 mM NaCl, 0.2% SDS). Reverse crosslinking was performed by adding 1 μl 20mg/ml proteinase K and incubating samples at 65 °C overnight. DNA was purified using the MinElute reaction cleanup kit following the manufacturer’s protocol. The purified DNA was end-repaired using an End-It DNA-Repair kit (Epicenter, Cat#ER81050) and added an A base at the 3’ end using the Klenow fragment (3’→5’ exo-) (NEB, Cat#M0212L). The DNA was then ligated with Y-shaped adaptors and amplified for 16 cycles using indexing primers with the PCR condition: 98 °C,10”; 67°C, 30”; 72°C, 30” as previously reported (*62, 63*). PCR products between 150-350 bp were isolated for sequencing on Illumina NovaSeq.

### Peak calling algorithm of cLoops2

Candidate peak regions were obtained from blockDBSCAN with PETs distance < 1 kb. If multiple parameters of *eps* and *minPts* were assigned, then blockDBSCAN with combinations of *eps* and *minPts* were performed, and candidate peaks were collected. For each candidate peak, the Poisson test was used to determine the reads’ statistical significance in the peak compared to nearby regions and the same region in the control sample (input or IgG).

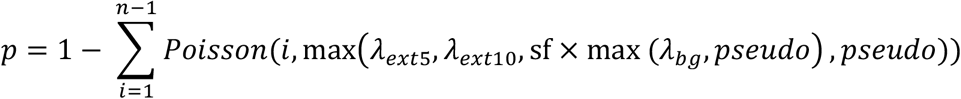

In the above formula, *n* is the observed PETs number in the candidate peak region from the test sample; *λ*_*ext*5_ and *λ*_*ext*10_ are the mean values of observed PETs number for the upstream and downstream same size 5 folds and 10 folds windows of the candidate peak region from the test sample, and insensitive mode, they are the median values; sf is the scaling factor between the control sample and the test sample, if no control sample assigned, sf = 0; otherwise sf is either library sequence depth ratio or the coefficient from linearly fitting of all candidate peak regions PETs number between test sample and control sample without intercepting; by default coefficient from linearly fitting is used as a scaling factor and in sensitive mode, the smaller scaling factor is used; *λ*_*bg*_ is the observed PETs number in the candidate peak region from the control sample; *pseudo* is used to control empirical background noise as an adjustable parameter, by default is 1.

The candidate peak with the highest RPKM value was reported for overlapped candidate peaks from multiple rounds of clustering results if the Poisson *P*-value was significant. Finally, all p-values were corrected by Bonferroni correction, and by default, corrected *p* < 0.01 was used to determine significant peaks. The algorithm is summarized as the cLoops2 callPeaks module.

### ChIC-seq analysis

Raw paired-end reads in FASTQ format were mapped to human reference genome hg38 by Bowtie2 (*64*). Mapped unique paired-end reads with MAPQ ≥ 10 were used for the following analysis. Peaks called by cLoops2 were performed with key parameters of cLoops2 callPeaks -eps 100,200 -minPts 5 -bdg IgG. Default parameters were used for peak calling by MACS2 (v2.2.6) except for H3K27ac ChIC-seq, for which –board was used. Peak calling by the SICER algorithm was performed by a faster implementation epic2 (v0.0.41) (*65*) with default parameters. Peak calling by HOMER was performed by findPeaks command, for CTCF ChIC-seq -style factor option was used and for H3K4me3 and H3K27ac ChIC-seq -style histone were assigned. Peak calling by SEACR were perfomred by both stringent mode and relaxed mode.

### Comparison of peaks called by various algorithms with ENCODE peaks

To use ENCODE peaks as a reference dataset for comparing peak-calling algorithms, ENCODE peaks were quantified first in ChIC-seq data (CUT&RUN or CUT&TAG for that part of the analyses). Only the ENCODE peaks that had signal densities of 2-fold higher than flanking upstream and downstream same-sized regions and 2-fold higher than the signals in the same region in the IgG (or input) control samples were kept for the analysis. Let *N*_*i*_ be the number of called peaks for a tool *i* for one factor, *n*_*i*_ be the number of called peaks overlapped with ENCODE reference peaks, *M* is the number of ENCODE reference peaks, and *m*_*i*_ is the number of ENCODE reference peaks overlapped with called peaks. Then *Precision*_*i*_ = *n*_*i*_/*N*_*i*_, *Sensitivity*_*i*_ = *m*_*i*_/*M*_*i*_ and 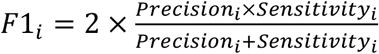. Overlapped peaks were obtained by BEDtools intersectBed command with -f option assigned 0.5 or 0 by requiring at least half of peaks were overlapped or minimum 1 bp.

### Loop calling algorithm of cLoops2

Candidate loops were obtained from blockDBSCAN. If multiple parameters of *eps* and *minPts* were assigned, then blockDBSCAN with combinations of *eps* and *minPts* were performed, and candidate loops were collected. The hypergeometric test and the Poisson test were the same as the description in cLoops (*28*). The binomial test for each candidate loop was updated as following,

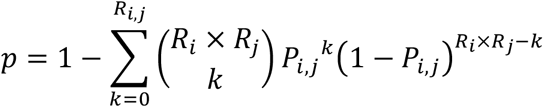

Where *R*_*i,j*_ is the number of PETs linking the candidate anchors *i*, *j*. *R*_*i*_ is the number of PETs located in candidate anchor i, and *R*_*j*_ is the number of PETs located in candidate anchor j. *P*_*i*,*j*_ is the possibility of observing 1 PET link of the two regions, estimated from local permutated background regions as 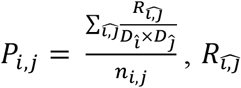 is the number of PETs linking the local permutated regions, 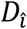 or 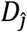 is the number of PETs in the permutated anchor 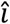 or 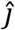, *n*_*i,j*_ is the total number of permutated loops. The changes generated similar trend binomial *P*-values with the previous method but removed the total number of PETs from the math formula.

### Aggregation analysis of loops

An 11 × 11 contact matrix was constructed for a loop from interacting PETs, together with its upstream and downstream same size 5 windows. An individual enrichment score for a loop was calculated as the number of PETs in the matrix center divided by all others’ mean values. The global enrichment score was the mean value of all enrichment scores for individual loops. Heatmap was plotted of the average matrix for all normalized 11 × 11 matrices. In contrast to the aggregation analysis of loops in Juicer (*48*), which by default filters some close loops before aggregation, cLoops2 runs the analysis for all loops by default.

### TrAC-looping data processing

Raw FASTQ files were processed by tracPre.py in the cLoops2 package mapped to human genome hg38 into BEDPE files. After processing into cLoops2 data by the cLoops2 pre module, cLoops2 samplePETs were used to do sub-sampling for the two samples into 100 million PETs for a fair comparison of loop calling and differentially enriched loop calling. Loops were called by cLoops2 callLoops module with parameters settings of -eps 200,500,1000 -minPts 5 -cut 1000. The cLoops2 callDiffLoops module called differentially enriched loops with default parameters settings. Raw reads of RNA-seq data were mapped to hg38 by STAR (v2.7.3a) (*66*), and fold changes of activated to resting CD4+ cells were obtained by Cuffdiff (v2.2.1) (*67*).

### Public data used

Re-analyzed public datasets of raw FASTQ files, mapped reads BED files, and ENCODE peaks were summarized in **Supplemental Table 1**.

### Data and code availability

Sequencing data generated in this study have been deposited to the Gene Expression Omnibus database with the accession of GSE179010. Results generated by cLoops2 in this study can be found in https://github.com/YaqiangCao/cLoops2_supp. All code of cLoops2 is open-source and available at GitHub https://github.com/YaqiangCao/cLoops2.

## FUNDING

This work was supported by the Intramural Research Program of the National Heart, Lung, and Blood Institute (KZ) and the 4DN Transformative Collaborative Project Award (A-0066) (KZ).

## ACKNOWLEDGEMENTS

We thank the NHLBI DNA sequencing facility for sequencing the NGS libraries. We thank Zhaoxiong Chen and Jing-Dong Jackie Han for essential discussion of the blockDBSCAN algorithm and Jonathan Perrie for cLoops2 documentation and testing help. This work utilized the computational resources of the NIH HPC Biowulf cluster (http://hpc.nih.gov). This research was supported by the Intramural Research Program of the National Institute of Health, National Heart, Lung and Blood Institute, NIH.

## AUTHOR CONTRIBUTIONS

YQC and KZ designed the project; YQC and ZXC implemented the blockDBSCAN algorithm; YQC finished all code, testing, documentation, and analysis. SL and QST carried out the Hi-TrAC and TrAC-looping experiments, and GR carried out the ChIC-seq experiments. All authors contributed to data analysis, interpretation and wrote the paper.

## Notes

### Competing Interest Statement

The authors have declared no competing interest.

