## Supplemental Information for "cLoops2: a full-stack comprehensive analytical tool for chromatin interactions"

#### Public data re-analyzed

Used datasets were summarized as following**.** GM12878 and K562 Hi-TrAC data were obtained from ***Liu. et al***. NHGRI-EBI GWAS Catalog data (*1*) were downloaded from https://www.ebi.ac.uk/gwas/api/search/downloads/full on November 11, 2020.

| Accession | Data Type | Cell Type | Factor | Reference |
|---|---|---|---|---|
| ENCLB620YVM, ENCLB779ZWL, (ENCODE) | ChIA-PET (FASTQ) | K562 | RAD21 | (*2*) |
| GSM2705045, GSM2705044, GSM2705043, (GEO) | HiChIP (FASTQ) | K562 | H3K27ac | (*3*) |
| GSM765405 (GEO) | RNA-seq (FASTQ) | K562 | None | (*4*) |
| ENCLB784HEF, | ChIA-PET | GM12878 | RAD21 | (*2*) |

| | | | | |
|---|---|---|---|---|
| ENCLB535GER, (ENCODE) | (FASTQ) | | | |
| GSM2705042, GSM2705041, (GEO) | HiChIP (FASTQ) | GM12878 | H3K27ac | (*3*) |
| GSM758559 (GEO) | RNA-seq (FASTQ) | GM12878 | None | (*4*) |
| GSM1551550, GSM1551551, GSM1551552, GSM1551553, GSM1551554, GSM1551555, GSM1551556, GSM1551557, GSM1551558, GSM1551559, GSM1551560, GSM1551561, GSM1551562, GSM1551563, GSM1551564, GSM1551565, GSM1551566, GSM1551567, GSM1551568, | Hi-C (FASTQ) | GM12878 | None | (*5*) |

| | | | | |
|---|---|---|---|---|
| GSM1551569, GSM1551570, GSM1551571, GSM1551572, GSM1551573, GSM1551574, GSM1551575, GSM1551576, GSM1551577, GSM1551578, GSM1551579, GSM1551580, GSM1551581, GSM1551582, GSM1551592, GSM1551592, GSM1551593, GSM1551594, GSM1551595, GSM1551596, GSM1551597, GSM1551598, (GEO) | | | | |
| GSM1872886 (GEO) | ChIA-PET (FASTQ) | GM12878 | CTCF | (*6*) |
| GSM2326178 | Trac-looping | Resting CD4+ | None | (*7*) |

| | | | | |
|---|---|---|---|---|
| GSM2326179<br>GSM2326180<br>GSM2782295<br>GSM2782296<br>(GEO) | (FASTQ) | | | |
| GSM2326181<br>GSM2326182<br>GSM2326183<br>(GEO) | Trac-looping<br>(FASTQ) | Activated CD4+ | None | (*7*) |
| GSM2326184<br>GSM2326185<br>(GEO) | RNA-seq | Resting CD4+ | None | (*7*) |
| GSM2326186<br>GSM2326187<br>(GEO) | RNA-seq | Activated CD4+ | None | (*7*) |
| ENCFF356LIU<br>(ENCODE) | ChIP-seq<br>(peaks) | GM12878 | CTCF | (*8*) |
| ENCFF367KIF<br>(ENCODE) | ChIP-seq<br>(peaks) | GM12878 | H3K27ac | (*8*) |
| ENCFF228GWY<br>(ENCODE) | ChIP-seq<br>(peaks) | GM12878 | H3K4me3 | (*8*) |
| ENCFF002DDJ<br>(ENCODE) | ChIP-seq<br>(peaks) | K562 | CTCF | (*8*) |
| ENCFF044JNJ<br>(ENCODE) | ChIP-seq<br>(peaks) | K562 | H3K27ac | (*8*) |
| ENCFF322IFF | ChIP-seq | K562 | H3K27me3 | (*8*) |

| | | | | |
|---|---|---|---|---|
| (ENCODE) | (peaks) | | | |
| ENCFF624XRN (ENCODE) | ChIP-seq (peaks) | K562 | H2A.Z | (*8*) |
| ENCFF183UQD (ENCODE) | ChIP-seq (peaks) | K562 | H3K4me1 | (*8*) |
| GSM3609747 (GEO) | CUT&RUN (aligned reads BED format) | K562 | CTCF | (*9*) |
| GSM3609748 (GEO) | CUT&RUN (aligned reads BED format) | K562 | CTCF | (*9*) |
| GSM3609749 (GEO) | CUT&RUN (aligned reads BED format) | K562 | CTCF | (*9*) |
| GSM3609750 (GEO) | CUT&RUN (aligned reads BED format) | K562 | CTCF | (*9*) |
| GSM3609755 (GEO) | CUT&RUN (aligned reads BED format) | K562 | H3K27ac | (*9*) |
| GSM3609756 (GEO) | CUT&RUN (aligned reads BED format) | K562 | H3K27ac | (*9*) |
| GSM3609757 (GEO) | CUT&RUN (aligned reads BED format) | K562 | H3K27ac | (*9*) |
| GSM3609758 (GEO) | CUT&RUN (aligned reads BED format) | K562 | H3K27ac | (*9*) |
| GSM3609759 | CUT&RUN | K562 | H3K27ac | (*9*) |

| | | | | |
|---|---|---|---|---|
| (GEO) | (aligned reads BED format) | | | |
| GSM3770938 (GEO) | CUT&RUN (aligned reads BED format) | K562 | H3K27me3 | (*9*) |
| GSM3770939 (GEO) | CUT&RUN (aligned reads BED format) | K562 | H3K27me3 | (*9*) |
| GSM3770940 (GEO) | CUT&RUN (aligned reads BED format) | K562 | H3K27me3 | (*9*) |
| GSM3609773 (GEO) | CUT&RUN (aligned reads BED format) | K562 | IgG | (*9*) |
| GSM3560260 (GEO) | CUT&Tag (aligned reads BED format) | K562 | H2A.Z | (*10*) |
| GSM3536516 (GEO) | CUT&Tag (aligned reads BED format) | K562 | H3K4me1 | (*10*) |
| GSM3536514 (GEO) | CUT&Tag (aligned reads BED format) | K562 | H3K27ac | (*10*) |
| GSM3560264 (GEO) | CUT&Tag (aligned reads BED format) | K562 | IgG | (*10*) |

**Supplemental Methods**

**Pre-processing of CUT&RUN and CUT&TAG data**

Pre-processed BED files mapped to hg19 were downloaded from GEO and converted to hg38 by liftOver for all following analyses. The same parameters were set for peak calling by other tools as described in the part on the comparison of ChIC-seq peak-callings. cLoops2 callPeaks with settings of -eps 100,200 -minPts 5 -sen for CTCF and H3K27ac, -eps 1000, 2000 -minPts 5 -sen for H3K27me3 were used to call peaks from CUT&RUN data. cLoops2 callPeaks with settings of -eps 100,200 -minPts 5 -sen for H2A.Z and H3K27ac, -eps 1000,2000 -minPts 5 -sen for H3K4me1, were used to call peaks from CUT&TAG data.

**Processing of Hi-C, HiChIP, and ChIA-PET data**

Pre-processing of Hi-C, H3K27ac HiChIP data, and RAD21 ChIA-PET data from raw FASTQ files to mapped BEDPE files was described as ***Liu et al***.

Sub-sampling of 150 million PETs from H3K27ac HiChIP data were used for all downstream analyses. H3K27ac HiChIP loops were called by cLoops2 callLoops module with parameters settings of -eps 200,500,1000 -minPts 20 -w -j -cut 10000. The cLoops2 callDiffLoops module called differentially enriched loops with default parameters.

Sub-sampling of 17 million PETs from RAD21 ChIA-PET data were used for all downstream analyses. RAD21 ChIA-PET loops were called by cLoops2 callLoops module with parameters settings of -eps 500,1000,2000 -minPts 3,5. The cLoops2 callDiffLoops module called differentially enriched loops with default parameters.

**Calling domains from Hi-TrAC data**

Domains were called from Hi-TrAC data based on a variant of insulation score (*11*), termed segregation score. A sliding window with fixed size (as a parameter) around each bin was used to obtain the sub matrix for calculating the segregation score for each bin. The contact matrix is further $\log_2$ transformed and calculated as a correlation matrix. For the up-right corner sub-matrix of the correlation matrix, all < 0 values are assigned as 0, and the mean value of the matrix is assigned as segregation score for the bin. After calculating segregation scores for all bins in one chromosome, z-score normalization was performed to the segregation scores. Finally, bins with >0 segregation scores were stitched together as candidate domains. This algorithm is implemented in cLoops2 callDomains module.

### Supplemental Figure 1

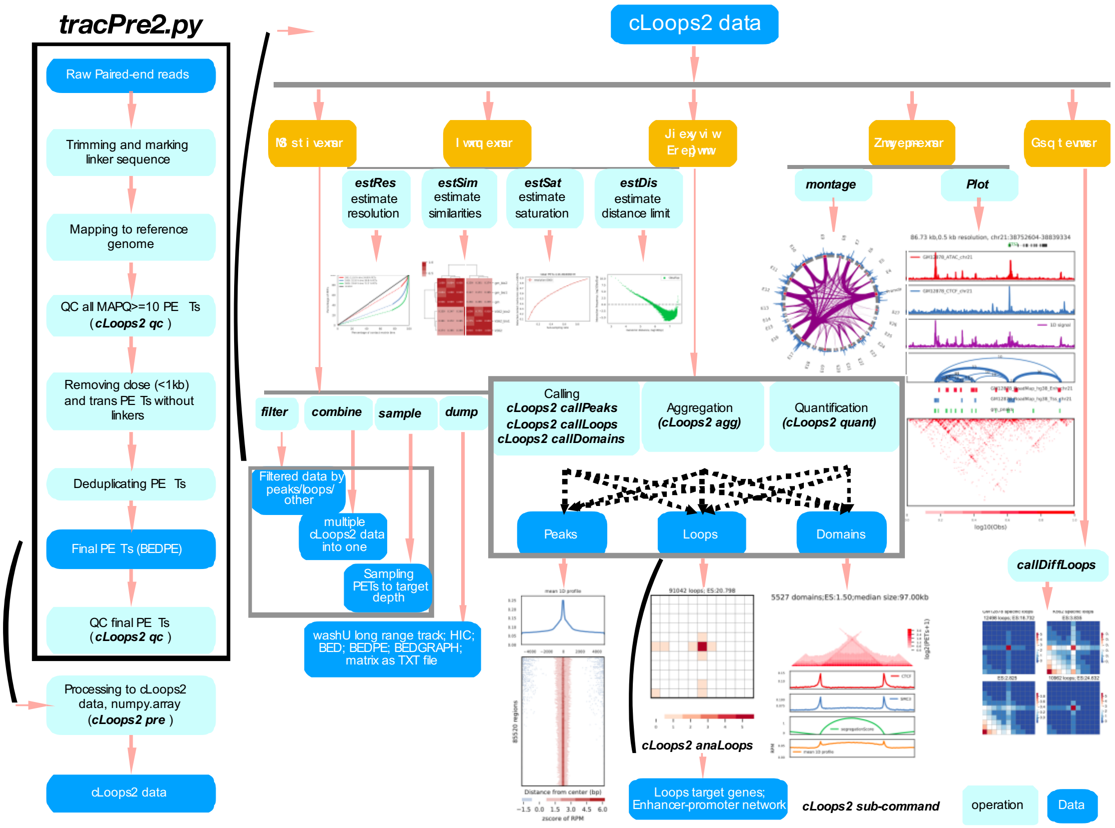


**Supplemental Figure 1. Overview of cLoops2 main modules architecture**

The main functions of cLoops2 were integrated into the main command cLoops2; meanwhile, extra analysis scripts were located in the script directory, such as Hi-TrAC data pre-processing analysis script tracPre2.py. In tracPre2.py, FASTQ files were first trimmed of linker sequence CTGTCTCTTATACACATCT for both ends. Only PETs with both ends length >=10bp were kept. Trimmed PETs were mapped to the target reference genome with Bowtie2 (*12*). Mapped PETs with MAPQ >=10 were converted to BEDPE files. PETs < 1kb without linker sequence in any end were further filtered. PCR replicates of PETs were filtered if the locations of both ends were identical. In cLoops2 main functions, modules were classified into 5 categories: IO operation, estimation, features analysis, visualization, and comparison. In IO operation category, cLoops2 pre module processes input BEDPE files into cLoops2 data; cLoops2 filterPETs module filters PETs by overlapping with peaks/loops or applies blockDBSCAN noise remove process; cLoop2 combine module combines multiple cLoops2 data directories; cLoops2 sample module does sampling PETs to target sequencing depth and cLoops2 dump module coverts cLoops2 data into other popular data formats such as BED, BEDPE, and HIC. In the estimation category, cLoops2 estRes module estimates interaction resolutions; cLoops2 estSim module estimate different datasets interaction similarities; cLoops2 estSat module estimates sequencing saturation for sequencing depth and cLoops2 estDis module estimates significant interaction distance limitations. The feature analysis category, features calling results from cLoops2 callPeaks module, callLoops module, and callDomains module can be further processed with cLoops2 agg module for aggregation analysis to draw global conclusions and cLoops2 quant module to obtain digital quantification for each feature to perform detailed analysis. An extra module called anaLoops was used to annotate loops target genes and obtain the potential enhancer-promoter interaction networks. In the visualization category, the cLoops2 montage module generates a Rehoboam plot for showing interactions for selected interest regions, and the cLoops2 plot module generates interactions heatmaps/arches/scatter plot together with genes genomic regions, loops, and other 1D profile data. In the comparison category, cLoops2 callDiffLoops performs differentially enriched loops analysis to find sample-specific loops, and the loops can be annotated using cLoops2 anaLoops to find potential specific target genes.

#### Supplemental Figure 2

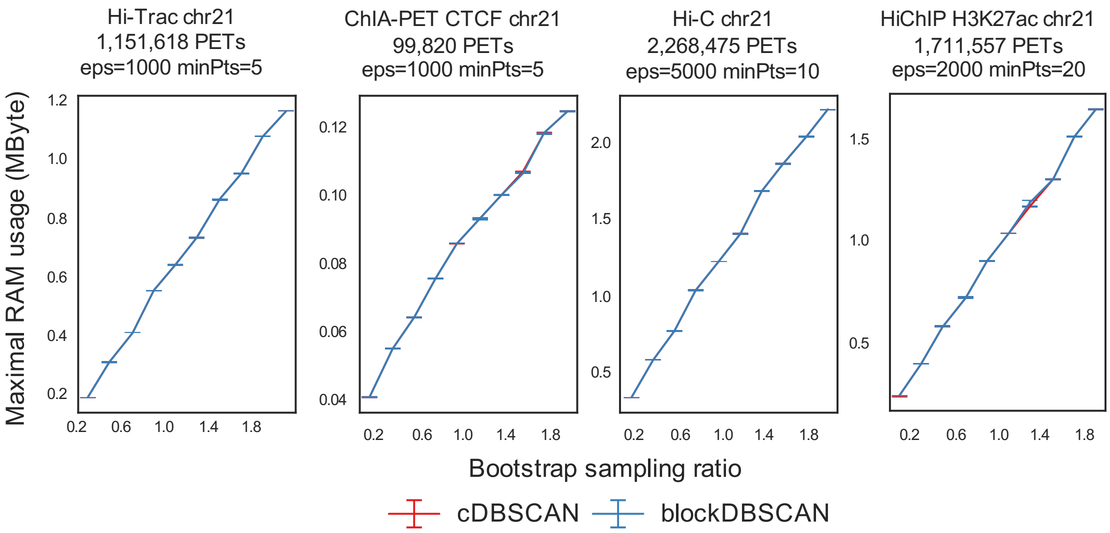


**Supplemental Figure 2. Comparison of memory usage between cDBSCAN and blockDBSCAN**

Comparison of memory usage using real interaction data of Hi-TrAC, CTCF ChIA-PET, Hi-C, and H3K27ac HiChIP data from GM12878 cells, based on 5 repeats sampling. The results from cDBSCAN and blockDBSCAN were highly similar; therefore, most parts of the curves were overlapped.

### Supplemental Figure 3

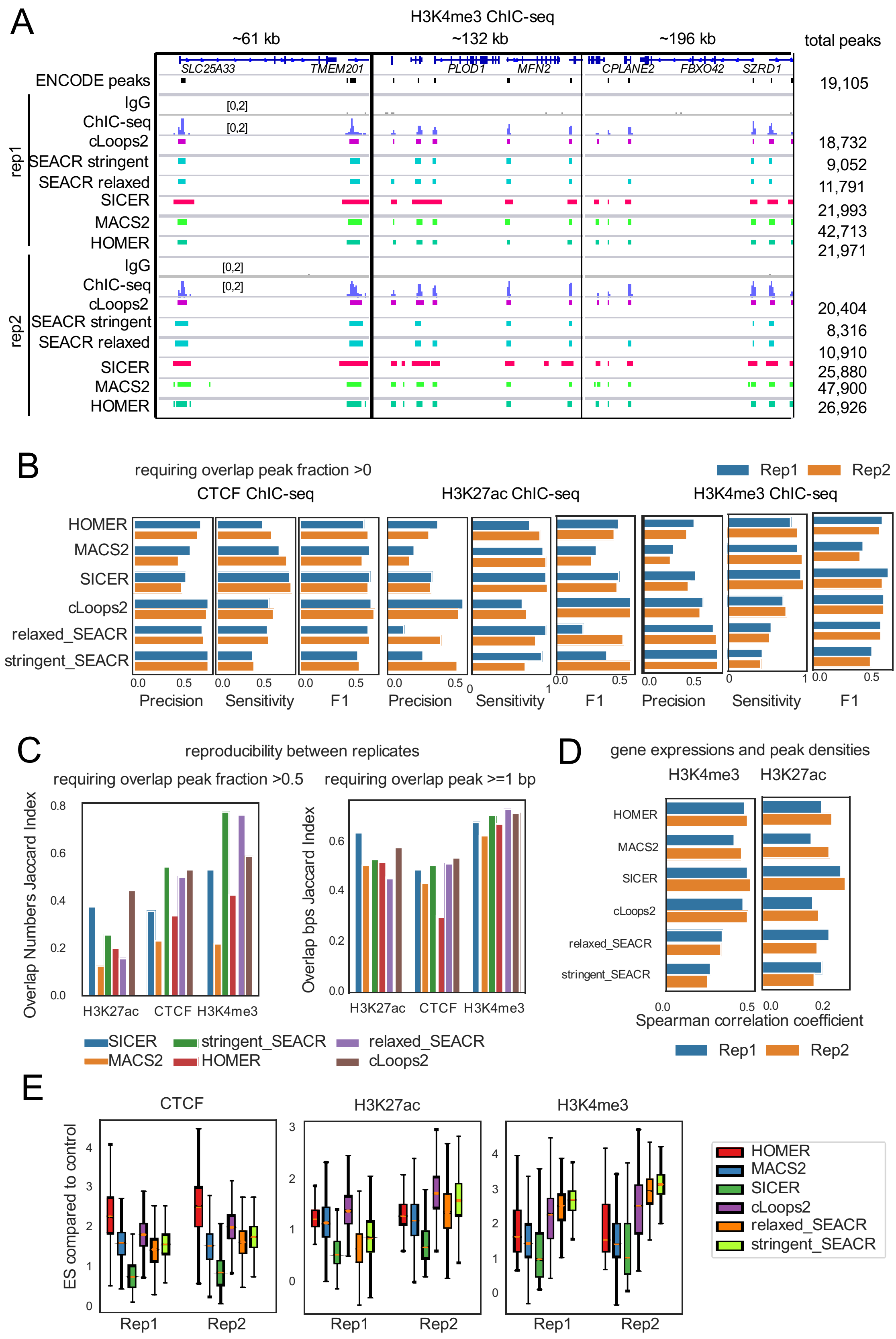


**Supplemental Figure 3. Comparison of ChIC-seq peak calling by various tools**

(A) Examples of H3K4me3 ChIC-seq profiles and peaks called by different tools.

(B) Precision, sensitivity, and F1 scores for peaks called by different tools comparing with ENCODE peaks, requiring peaks overlap >=1 bp.

(C) Reproducibility between ChIC-seq replicates. The left panel requiring peaks overlap fraction >=0.5, and the right panel requiring peaks overlap >=1bp.

(D) Correlation between gene expression and ChIC-seq peak densities.

(E) ChIC-seq signal enrichment score (ES) of called peak regions comparing to IgG control.

### Supplemental Figure 4

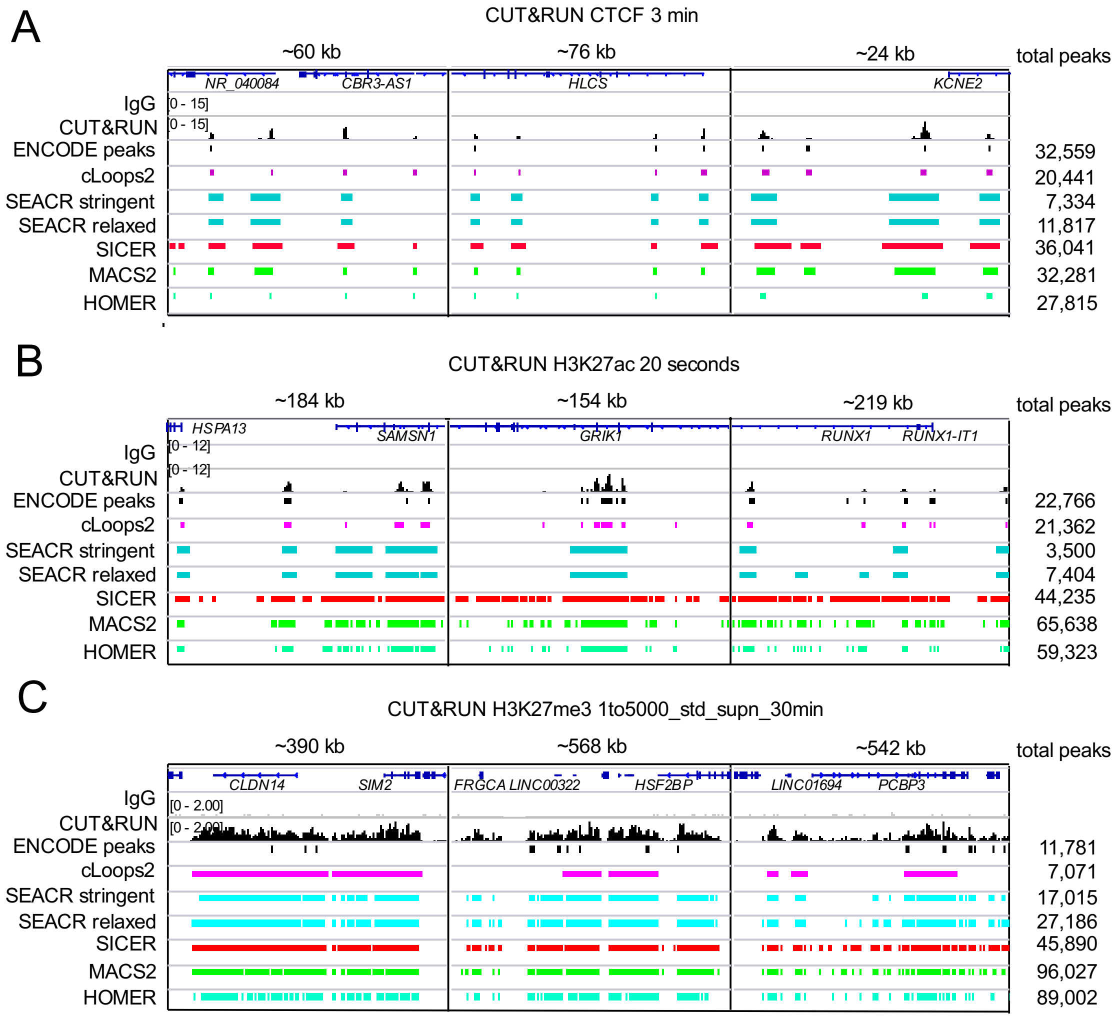


**Supplemental Figure 4. Examples of peaks called by cLoops2 and other tools from CUT&RUN data in K562 cells**

(A) Examples of CTCF CUT&RUN binding profiles and peaks called by different tools.

(B) Examples of H3K27ac CUT&RUN profiles and peaks called by different tools.

(C) Examples of H3K27me3 CUT&RUN profiles and peaks called by different tools.

### Supplemental Figure 5

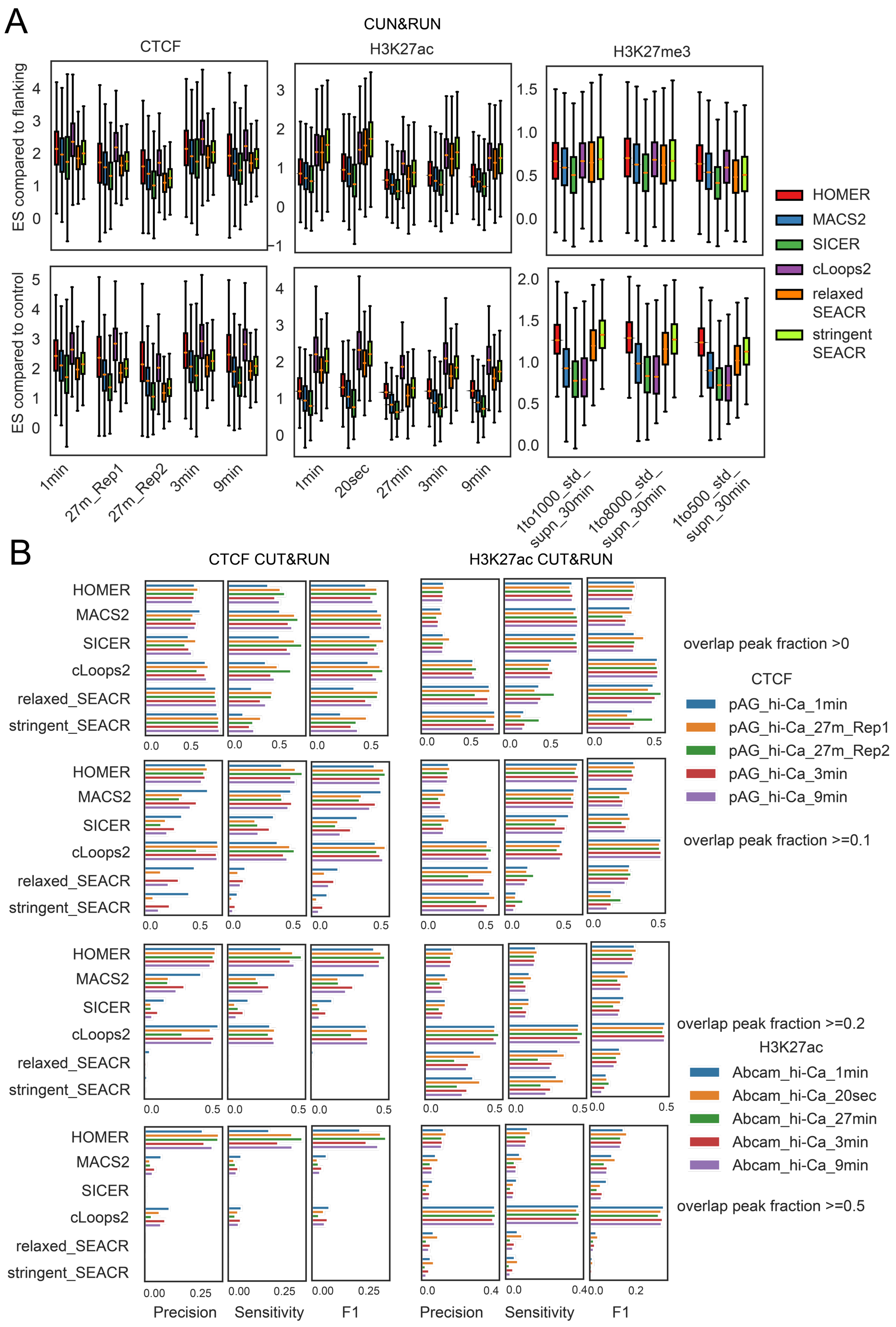


**Supplemental Figure 5. Global comparison of peaks called by cLoops2 and other tools from CUT&RUN data**

(A) The signal enrichment scores (ES) of peak regions compared to peak-flanking regions and IgG control. The peaks were called by cLoops2 and other tools from CTCF, H3K27ac and H3K27me3 CUT&RUN data.

(B) Precision, sensitivity, and F1 scores for peaks called by different tools in comparison with the ENCODE peaks.

### Supplemental Figure 6

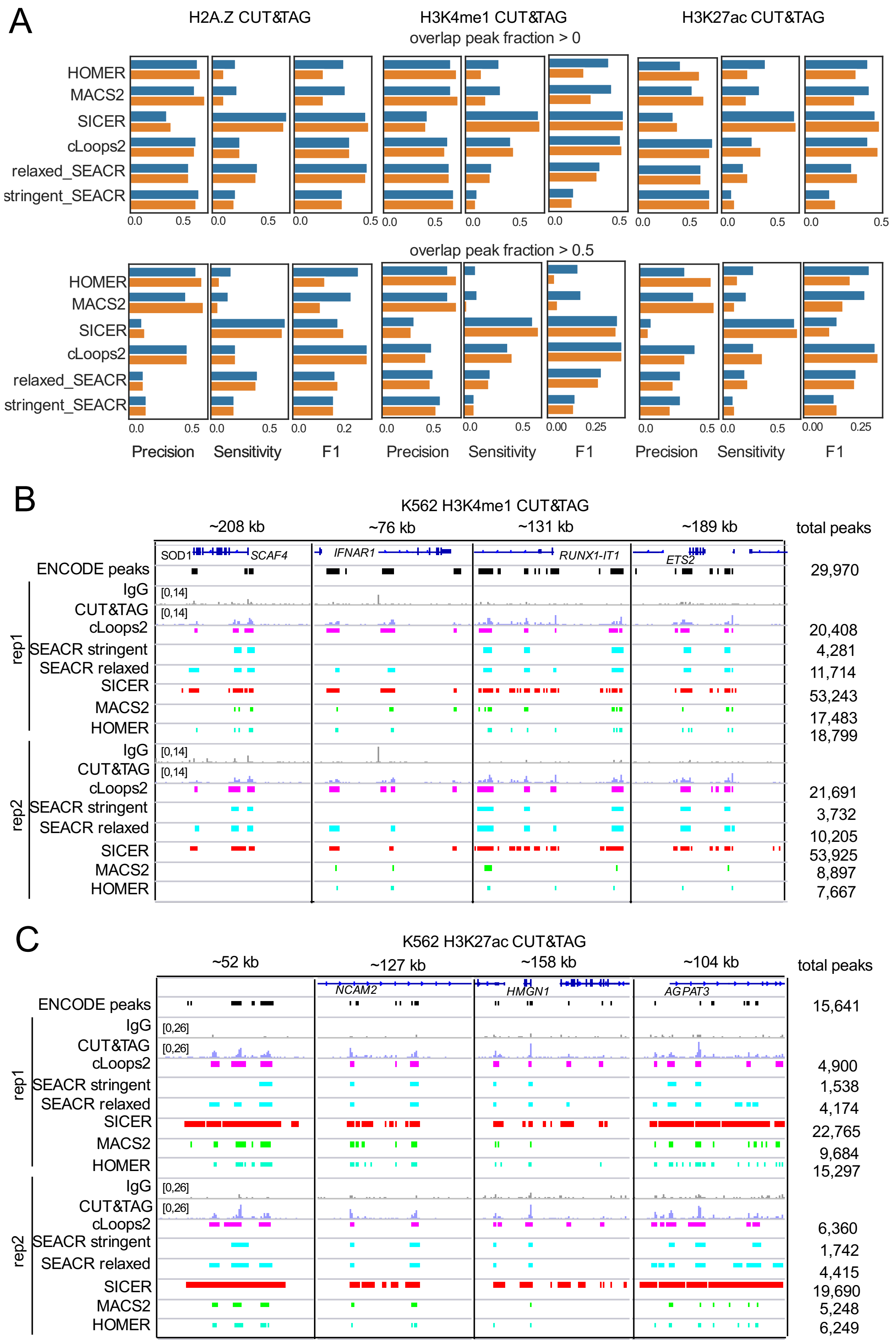


**Supplemental Figure 6. Global comparison of peaks called by cLoops2 and other tools from CUT&TAG data**

(A) Precision, sensitivity, and F1 scores for peaks called by different tools in comparison with the ENCODE peaks.

(B) Examples of H3K4me1 CUT&TAG profiles and peaks called by different tools.

(C) Examples of H3K27ac CUT&TAG profiles and peaks called by different tools.

#### Supplemental Figure 7

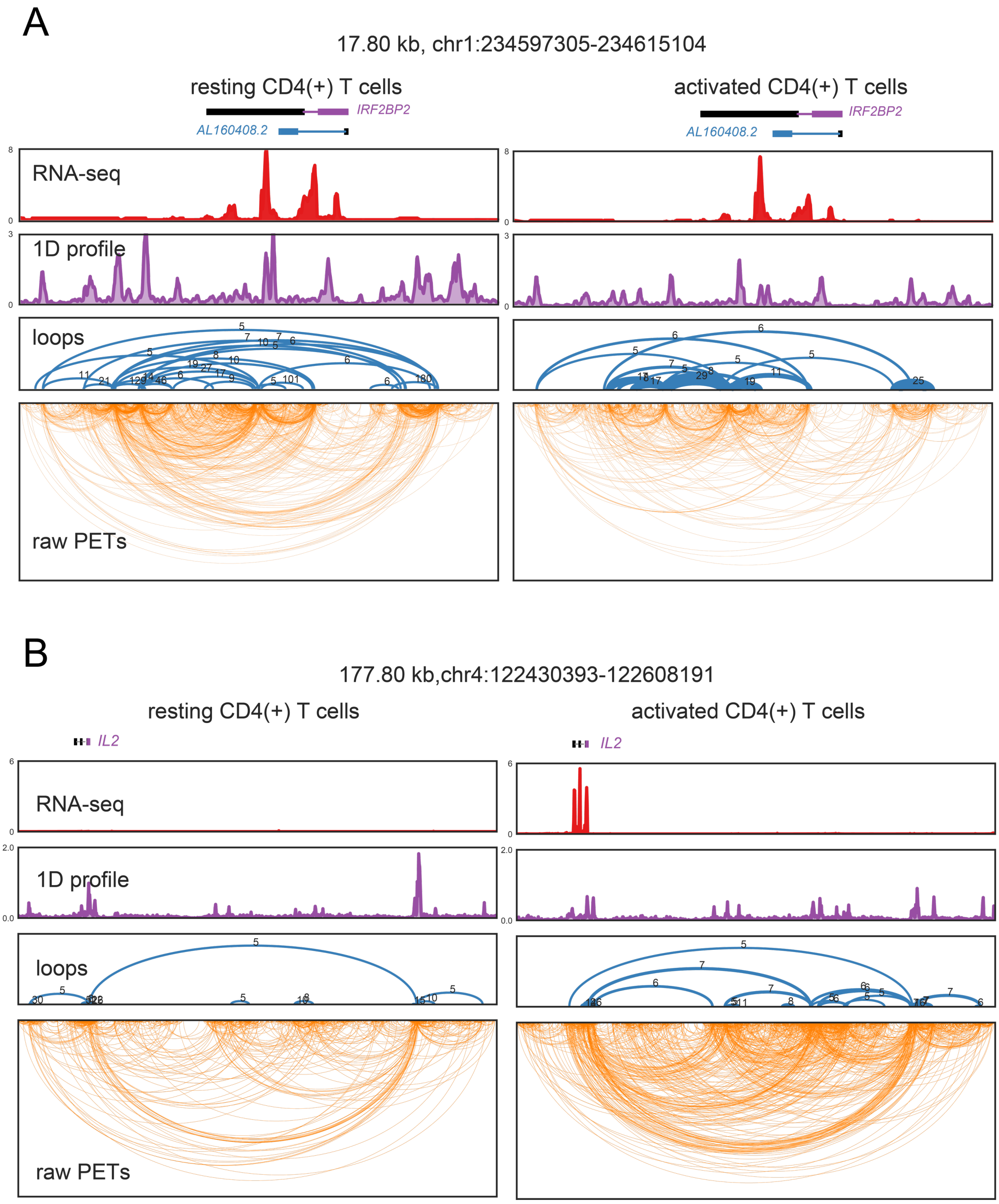


**Supplemental Figure 7. Examples of loops called by cLoops2 from CD4 T cell TrAC-looping data**

(A) Loops called by cLoops2 from TrAC-looping data around the *IRF2BP2* gene. The plots were generated by the cLoops2 plot module with -arch parameter. Blue color marked the first exon of the gene in the positive strand, and purple color marked the first exon of the gene in the negative strand.

(B) Loops called by cLoops2 from TrAC-looping data around the *IL2* gene. The plots were generated by the cLoops2 plot module with -arch parameter.

#### Supplemental Figure 8

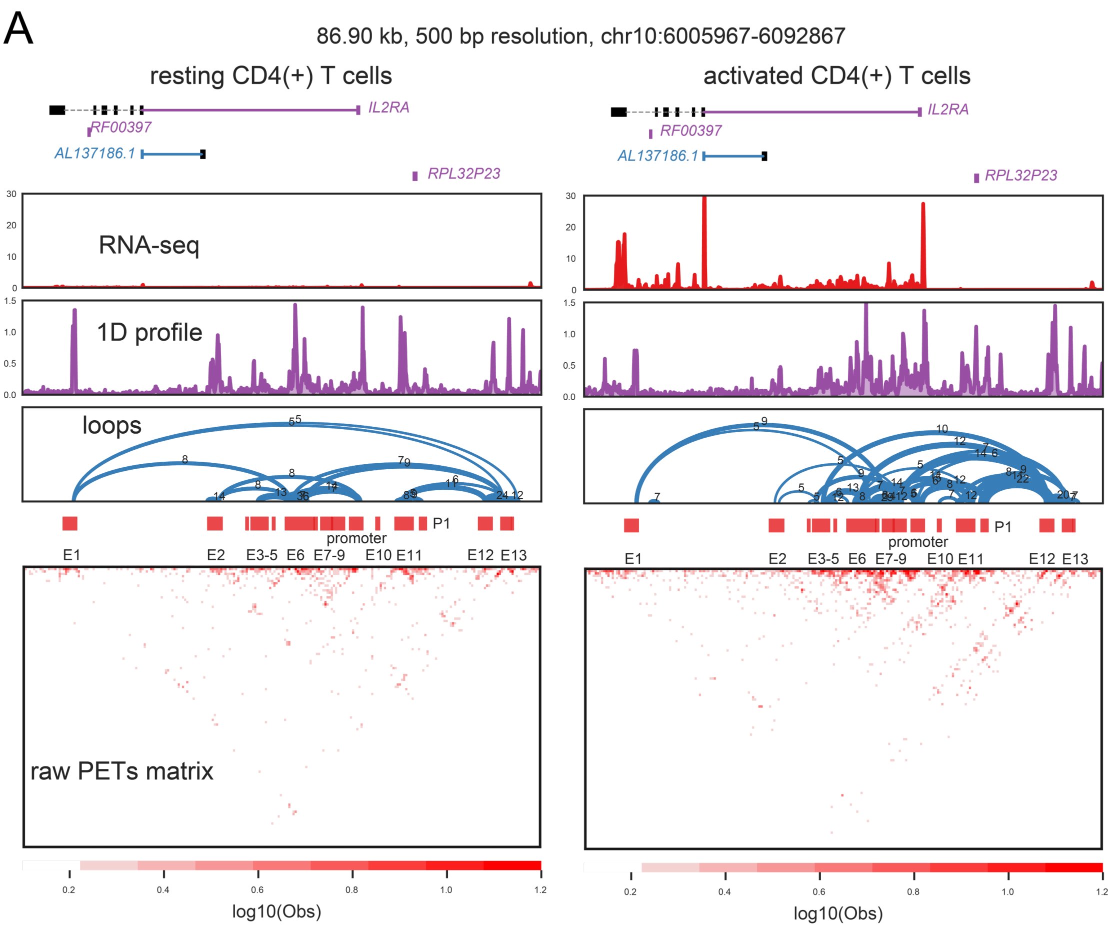


**Supplemental Figure 8. The loops called by cLoops2 around the *IL2RA* gene promoter in resting and activated CD4 T cells**

Supplemental Figure 9

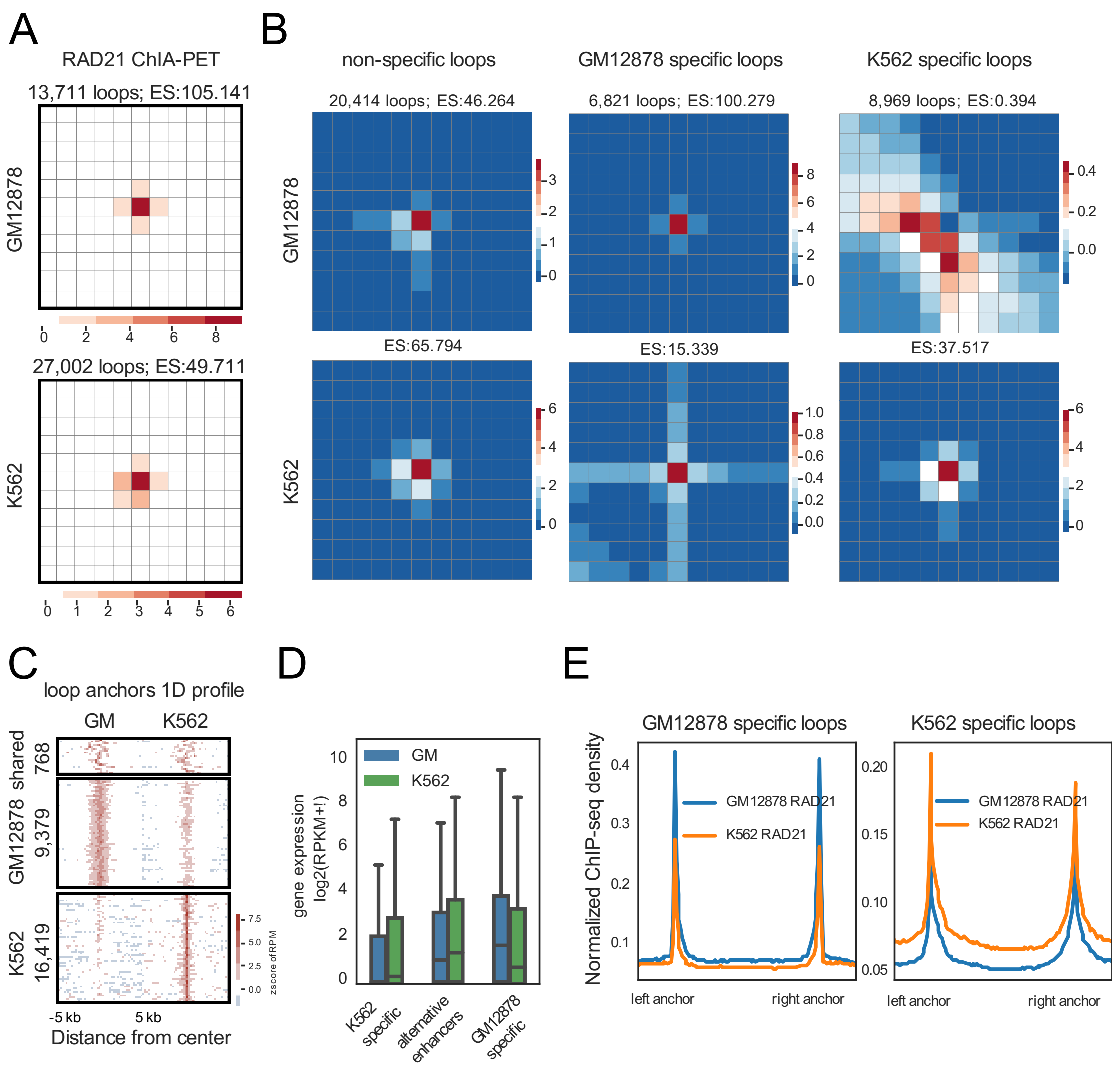


**Supplemental Figure 9. Differentially enriched loops from RAD21 ChIA-PET data called by cLoops2**

(A) Aggregation analysis of loops called by cLoops2 from the RAD21 ChIA-PET data from GM12878 and K562 cells.

(B) Aggregation analysis of the non-specific and cell-specific RAD21 guided loops between GM12878 and K562.

(C) Aggregation analysis of ChIA-PET 1D signals of the combined anchors from the non-specific and cell-specific loops.

(D) Gene expression association with cell-specific RAD21 guided loops. Alternative enhancers referred to the genes whose promoters were looped to different enhancers in the two different cell types.

(E) Aggregated RAD21 ChIP-seq signals on cell-specific RAD21-guided loops. ChIP-seq data were obtained from ENCODE.

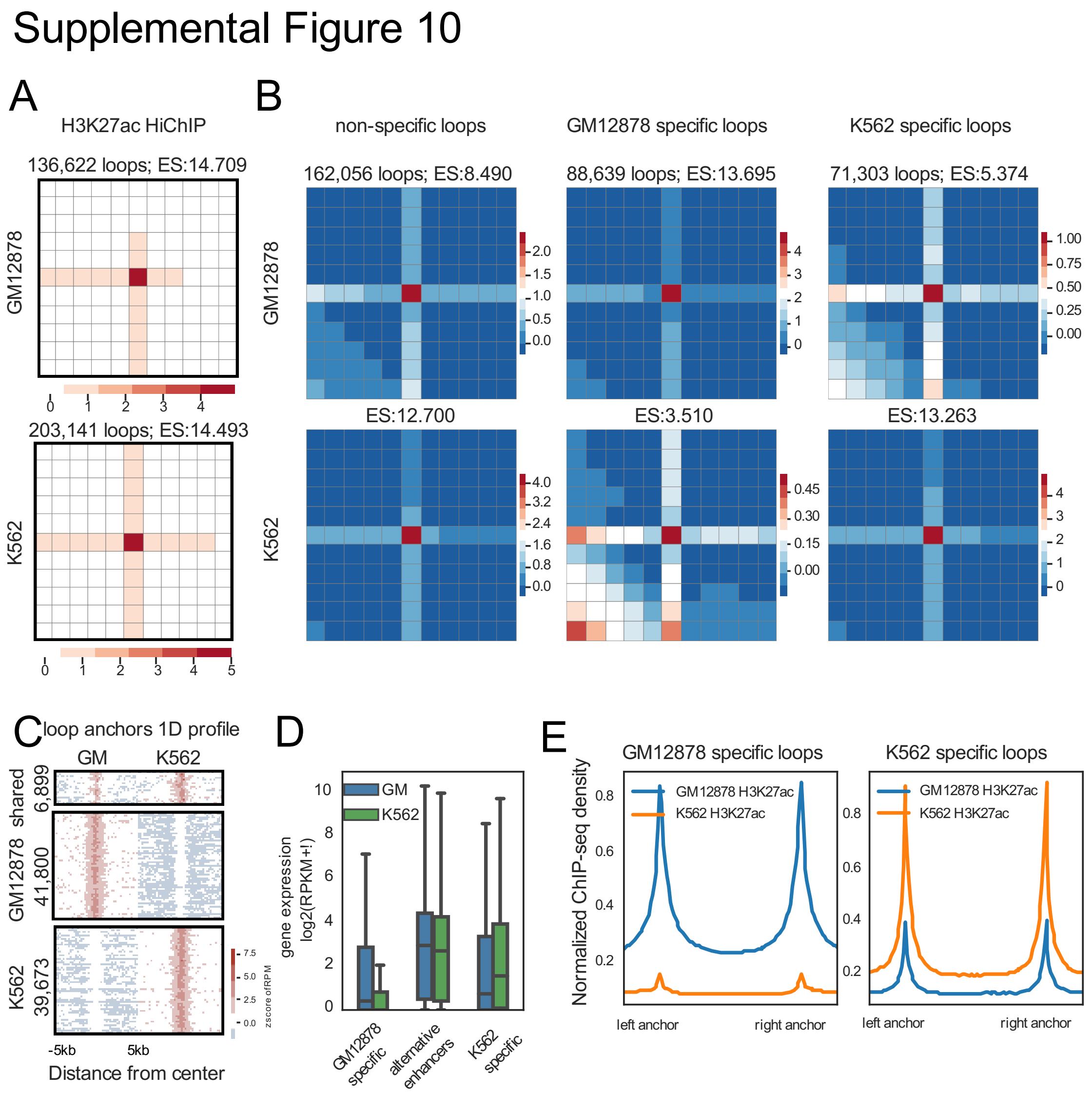


**Supplemental Figure 10. Differentially enriched loops from H3K27ac HiChIP data called by cLoops2**

(A) Aggregation analysis of loops called by cLoops2 from the H3K27ac HiChIP data from GM12878 and K562 cells.

(B) Aggregation analysis of the non-specific and cell-specific H3K27ac associated loops between GM12878 and K562.

(C) Aggregation analysis of HiChIP 1D signals of the combined anchors from the non-specific and cell-specific loops.

(D) Gene expression is associated with cell-specific H3K27ac associated loops.

(F) Aggregated H3K27ac ChIP-seq signals on the cell-specific H3K27ac-associated loops. ChIP-seq data were obtained from ENCODE.

#### Supplemental Figure 11

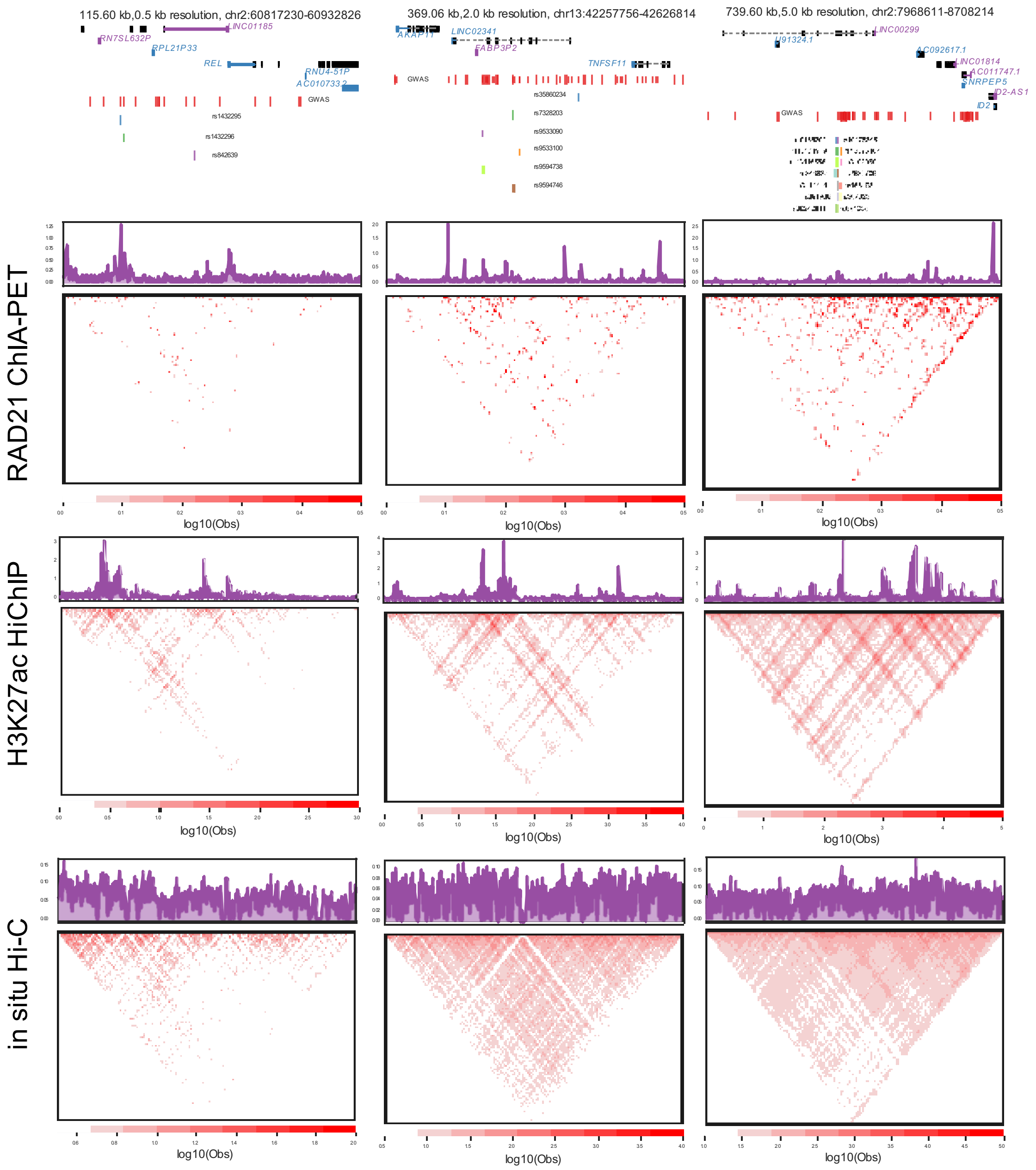


**Supplemental Figure 11. GM12878 RAD21 ChIA-PET, H3K27ac HiChIP, and Hi-C data for the three lncRNAs shown in Figure 5C-E**
